# LRP10 as a novel α-synuclein regulator in Lewy body diseases

**DOI:** 10.1101/2023.05.12.540510

**Authors:** Ana Carreras Mascaro, Martyna M. Grochowska, Valerie Boumeester, Natasja F. J. Dits, Ece Naz Bilgiҫ, Guido J. Breedveld, Leonie Vergouw, Frank Jan de Jong, Martin E. van Royen, Vincenzo Bonifati, Wim Mandemakers

## Abstract

Autosomal dominant variants in *LRP10* have been identified in patients with Lewy body diseases (LBDs), including Parkinson’s disease (PD), Parkinson’s disease-dementia (PDD), and dementia with Lewy bodies (DLB). Nevertheless, there is little mechanistic insight into the role of LRP10 in disease pathogenesis. In the brains of non-demented individuals, LRP10 is typically expressed in non-neuronal cells like astrocytes and neurovasculature, but in idiopathic and genetic cases of PD, PDD, and DLB it is also present in α-synuclein-positive neuronal Lewy bodies. These observations raise the questions of what leads to the accumulation of LRP10 in Lewy bodies and whether a possible interaction between LRP10 and α-synuclein plays a role in disease pathogenesis. Here, we demonstrate that wild-type LRP10 is secreted via extracellular vesicles (EVs) and can be internalised via clathrin-dependent endocytosis. Additionally, we show that LRP10 secretion is highly sensitive to autophagy inhibition, which induces the formation of atypical LRP10 vesicular structures in neurons in human induced pluripotent stem cells (iPSC)-derived midbrain-like organoids (hMLOs). Furthermore, we show that LRP10 overexpression leads to a strong induction of monomeric α-synuclein secretion, together with time-dependent, stress-sensitive changes in intracellular α-synuclein levels. Interestingly, patient-derived astrocytes carrying the *c.1424+5G>A LRP10* variant secrete aberrant high-molecular-weight species of LRP10 in EV-free media fractions. Finally, we show that the truncated LRP10^splice^ protein binds to wild-type LRP10, reduces LRP10 wild-type levels, and antagonises the regulatory effect of LRP10 on α-synuclein levels and distribution. Together, this work provides initial evidence for a functional role of LRP10 in LBDs by regulating intra- and extracellular α-synuclein levels, and pathogenic mechanisms linked to the disease-associated *c.1424+5G>A LRP10* variant, pointing towards potentially important disease mechanisms in LBDs.

## Introduction

Lewy body diseases (LBDs) are neurodegenerative disorders characterised by the widespread presence of insoluble α-synuclein-containing aggregates, accompanied by the loss of dopaminergic neurons in the substantia nigra pars compacta (1). The main LBDs include Parkinson’s disease (PD), Parkinson’s disease-dementia (PDD), and dementia with Lewy bodies (DLB), which collectively affect over 8 million people worldwide (2, 3). PD, PDD, and DLB display overlapping clinical features that can include parkinsonian motor signs and cognitive impairment. The main distinction in their diagnosis is based on the temporal sequence of the onset of these symptoms. In addition to sharing many clinical and neuropathological features, LBDs also present shared genetic factors, such as *GBA1*, *SNCA*, and *LRP10* genetic variants (4–9).

The low-density lipoprotein receptor (LDLR)-related protein 10 (LRP10) is a single-pass transmembrane protein and a member of the LDLR family (10). LRP10 function has been linked to the intracellular vesicle transport pathway due to its intracellular localisation pattern and interaction with clathrin adaptors and the retromer complex (9, 11–15). Increasing genetic and neuropathological evidence has positioned LRP10 as a novel player in LBDs. Several studies have identified potentially pathogenic variants in the encoding gene *LRP10* in autosomal dominant forms of PD, PDD, and DLB (9, 16–23). Furthermore, recent work from Grochowska et al. (11) showed that 71 to 95% of mature brainstem Lewy bodies in ten patients suffering from idiopathic and genetic cases of PD, PDD, and DLB were strongly positive for LRP10, even though its expression was almost limited to astrocytes and neurovasculature in non-demented control brains. Besides LBDs, *LRP10* variants were also identified in patients with progressive supranuclear palsy and amyotrophic lateral sclerosis (24, 25). Furthermore, LRP10 function has been associated with Alzheimer’s disease (AD). In addition to its physical interaction with SORL1, LRP10 has been described to be a receptor for the amyloid precursor protein (APP) and ApoE-carrying lipoproteins, and it has been shown to be a driver of an AD subtype (11, 26–28). Despite this effort, LRP10 function and its implication in neurodegeneration remain incompletely characterised.

The deposition of α-synuclein aggregates is a defining pathological feature of LBDs, and the cell-to-cell transmission of α-synuclein pathology is considered a major player in the disease progression (29). α-Synuclein is a presynaptic protein that is normally found in a soluble cytosolic state or bound to cellular membranes (30–33). However, several forms of α-synuclein have been associated with increased misfolding and aggregation potential, including the acquisition of a β- sheet amyloid conformation, post-translational modifications, and genetic variations (32, 34). The factors triggering the misfolding and deposition of α-synuclein in the human brain are still not clear, and they may vary per patient, affected brain area, and disease (35). In contrast, the transmission of soluble and pathological α-synuclein is well established, and it has been identified on the cellular level but also between anatomically-connected brain areas and between the brain and other organs, such as the gastrointestinal tract and peripheral muscles (36–43).

Based on these observations, we aimed to investigate whether the presence of LRP10 in Lewy bodies could be caused by the transfer of LRP10 from non-neuronal cells and subsequent internalisation by dopaminergic neurons, and whether LRP10 may play a role in regulating α- synuclein in LBD pathogenesis. Using human induced pluripotent stem cells (iPSC)-derived midbrain-like organoids (hMLOs), astrocytes, and LRP10-overexpressing cells as model systems, we found that LRP10 can be secreted via extracellular vesicles (EVs) and is highly sensitive to autophagy blockade by Bafilomycin A1 (BafA1), which induced the formation of atypical LRP10 structures in neurons in hMLOs. Additionally, we found that LRP10 overexpression induces changes in α-synuclein intracellular levels over time, and strongly promotes α-synuclein secretion via the endoplasmic reticulum (ER) and proteasomal pathways. Lastly, we show that the *LRP10 c.1424+5G>A* genetic variant found in LBD patients (9) affects LRP10 secretion and internalisation and has a dominant negative effect on LRP10-mediated regulation of intra- and extracellular α-synuclein levels. Altogether, our results show initial evidence for the role of LRP10 as an α-synuclein regulator in LBDs.

## Methods

### Cloning

To generate the pLVX-EF1α-LRP10-IRES-NeoR plasmid used for LRP10 overexpression, *LRP10* sequence-verified cDNA was subcloned from the pcDNA™3.1-LRP10-V5-His-TOPO® plasmid described previously (9) into the pLVX-EF1α-IRES-mCherry plasmid (Takara Bio, 631987). Briefly, LRP10 was inserted after the EF1α promotor in EcoRI and BamHI-digested pLVX-EF1α- IRES-mCherry using Gibson Assembly® (NEB) according to the manufacturer’s specifications. Next, the backbone was purified after MluI and MscI digestion from gel using the QIAquick® Gel Extraction Kit (Qiagen) excluding mCherry. The Neomycin resistance gene (NeoR) was subcloned from the pcDNA™3·1-LRP10-V5-His-TOPO® plasmid and located in place of mCherry via Gibson Assembly®. The negative control plasmid pLVX-EF1α-IRES-NeoR was generated by restriction digestion (EcoRI and BamHI), blunt end generation, and T4 ligation of pLVX-EF1α- LRP10-IRES-NeoR. The pLVX-EF1α-LRP10^splice^-IRES-NeoR plasmid was generated by PCR of specific regions of LRP10 from the pLVX-EF1α-LRP10-IRES-NeoR plasmid (LRP10^splice^ sequence can be found in Supplementary resource 1), and placed into the same plasmid after EcoRI, BamHI, BsmBI, and Xcm I restriction digestion to linearize the backbone and digest LRP10 wild-type, respectively.

The pcDNA™3.1-mCherry-LRP10 plasmid was generated from the pcDNA™3.1-LRP10-V5- His-TOPO® plasmid. Briefly, a stop codon was first introduced between LRP10 and V5-His sequences (described in detail in Quadri and Mandemakers, et al. (9)). Next, LRP10 was amplified by PCR introducing a BspEI restriction site at the N-terminal end of LRP10 and reintroduced into the same backbone by T4 ligation. Finally, mCherry was introduced into the pcDNA™3.1-BspEI-LRP10 plasmid via PCR amplification from the pmCherryN1 plasmid (Clontech) via the addition of BspEI restriction sites in the primer sequences, restriction digestion with BspEI, and T4 ligation.

To produce the plasmid used to generate LRP10 knock-out neuronal precursor cells (NPCs), pSpCas9-GFP-LRP10gRNA (11, 44) was digested with BseRI and BsrGI, and GFP was replaced with Puromycin resistance gene (PuroR, subcloned from GIPZ Non-silencing Lentiviral shRNA Control, Horizon, RHS4346) via Gibson Assembly®, resulting in the pSpCas9-PuroR- LRP10gRNA plasmid.

The pCMVsport6-SNCA (Horizon, clone Id: 6147966) plasmid was used for α-synuclein overexpression experiments.

To generate the doxycycline-inducible LRP10^splice^ cell line, the pCW57-LRP10^splice^-2A-PuroR was used. First, LRP10^splice^ was subcloned from the pLVX-EF1α-LRP10^splice^-IRES-NeoR plasmid via PCR and placed after the rtTA (reverse tetracycline-controlled transactivator)-Advanced promoter in the AgeI-digested pCW57-MCS1-2A-MCS2 (Addgene, 71782, from Adam Karpf) plasmid via Gibson Assembly®. All plasmids were verified by Sanger sequencing.

### HEK-293T and HuTu-80 cell culture and transfection

HEK-293T and HuTu-80 (ATCC®, HTB-40™) cell lines were expanded in growth medium (DMEM or DMEM:F12, respectively; Thermo Fisher Scientific, 10% fetal bovine serum (FBS), 1% Penicillin-Streptomycin) at 37 °C/5% CO^2^. Cells were split every two-three days with Trypsin–EDTA (Gibco). Prior to transfection, HEK-293T cells were seeded to a 60-70% confluency in tissue culture plates. Transfections were performed using the GeneJuice® transfection reagent (Merck) according to the manufacturer’s specifications, and the samples were further processed 48 h after transfection or when specified.

### Primary cell lines

Details regarding the primary cell lines used in this study can be found in Table 1. The Control-1 cell line was obtained from the Gladstone Institutes. The Control-3 line was commercially available (Gibco). The Control-2 cell line and the line derived from the DLB patient carrying the *LRP10 c.1424+5G>A* variant (9) were obtained within study protocols approved by the medical ethical committee of Erasmus MC and conformed to the principles of the Declaration of Helsinki. The participating subjects provided written informed consent for the use of the material for research purposes. The human embryonic stem cell lines (HUES) used in the RT-qPCR experiments (Supplementary Fig. 4) were a kind gift from Chad A. Cowan (Harvard University, Harvard Stem Cell Institute) (45).

**Table 1:** Details of the control and patient clonal lines included in this study.

| Donor | Source | Donor Age | Donor Sex | Donor Sampling Site | Derivation |
| --- | --- | --- | --- | --- | --- |
| Control-1 | WTC, Gladstone Institutes (Coriell Research Institute, GM25256) | 30 | Male | Dermal fibroblasts | hMLOs and astrocytes; LRP10 knock-out (LRP10<sup>KO</sup>) astrocyte line |
| Control-2 | *In house* | 57 | Male | Dermal fibroblasts | Astrocytes |
| Control-3 | Gibco™, Lot number: 18388 | 34 | Female | Dermal fibroblasts | hMLOs |
| DLB Patient | *In house* | 68 | Female | Dermal fibroblasts | Clones 1 and 2–astrocytes<br>Clone 3 - hMLOs |

### Generation and characterisation of human iPSCs

Control-1 was generated and characterised as previously described by Mandegar et al. (46). Control-2 was reprogrammed and characterised as described earlier (9, 47). Patient iPSCs were collected, and the total RNA was isolated using the ReliaPrep™ RNA MiniPrep kit (Promega) as recommended by the manufacturer. For each sample, on-membrane DNase I digestion was performed according to the manufacturer’s protocol (Promega). The integrity of the total RNA was assessed using spectrophotometric analysis using NanoDrop 2000/2000c. 260/280 values of ∼2.0 indicated pure RNA. Next, 2 µg of the total RNA was converted into cDNA using the SuperScript II Reverse Transcriptase kit (Invitrogen™). qPCR was performed on CFX384 Real-Time PCR Detection System (Bio-Rad) with ∼ 20 ng cDNA per reaction. The following cycling conditions were used: 3 min at 95 °C (initial denaturation), 40 cycles of 30 s at 95 °C, 30 s at 58 °C, and 72 s at 72 °C. The normalised expression of each target gene was calculated using the delta–delta Cq method. *GAPDH* was used as a reference gene. HUES 9 line was used for the normalisation of the gene expression. Primers were adapted from a previously published article (48).

### Generation of neural progenitor cells

The procedure to generate neural progenitor cells (NPCs) was adapted from Reinhardt et al. (49) and is described in detail in Grochowska et al. (11). Briefly, iPSCs were detached and cultured as embryoid bodies in suspension in a shaker. Neural induction was initiated by promoting Wnt and Shh signalling with the addition of SB431542 (Tocris, 1614), dorsomorphin (Abcam, ab120843), CHIR (Sigma-Aldrich, SML1046), and Purmorphamine (Stem Cell Technologies, 72202). Embryoid bodies showing neuroepithelial development were selected, dissociated, and plated on Matrigel® (Corning, 356238)-coated plates as NPCs. From the sixth passage, NPCs were kept in NPC medium, consisting of N2B27 medium (DMEM/F-12 – Neurobasal in 1:1 ratio, 1% B27 w/o Vitamin A, 0.5% N2, 1% Penicillin-Streptomycin, all from Thermo Fisher Scientific) supplemented with 3 μM CHIR, 200 μM ascorbic acid (AA, Sigma-Aldrich, A92902), and 0.5 μM Smoothened Agonist (SAG, Abcam, ab142160). Cells were kept on Matrigel-coated plates, refreshed every other day, and split with accutase (Sigma-Aldrich) in a 1:10-1:20 ratio.

### Astrocyte differentiation

NPCs were differentiated into astrocytes according to published protocols (11, 49) with minor modifications. First, 75.000 NPCs/cm^2^ were seeded on Matrigel® (Corning)-coated plates with N2B27 medium supplemented with 10 ng/mL FGF-basic (PeproTech, 100-18B) and 10 ng/mL EGF (PeproTech, AF-100-15). After two days, this medium was switched to N2 medium (Advanced DMEM/F-12, 4% FBS, 1% N2, 1% Penicillin-Streptomycin) supplemented with 10 ng/mL CNTF (PeproTech, 450-13). After 12 days, cells were grown in N2 medium with 10 ng/mL EGF. At 2 months of differentiation, cells were frozen in N2 medium with 10% DMSO. All the lines were thawed simultaneously and kept in N2 medium with 10 ng/mL EGF for 2-3 weeks, followed by at least 10 days of N2 medium with 10 ng/mL CNTF. Cells were refreshed every other day. One week before terminating cultures, cells were treated with 100 μM dbcAMP in N2 medium. Cells were cultured for at least 3 months. Astrocyte cultures were split with accutase (Sigma-Aldrich) at a 1:2-1:5 ratio when maximal density was reached, leaving most cell clumps intact.

### hMLOs differentiation

NPCs were differentiated into hMLOs according to published protocols (50, 51) with minor modifications. NPCs were dissociated with accutase (Sigma-Aldrich), briefly centrifuged, and resuspended in NPC medium. Cells were counted using the TC20 automated cell counter (Bio-Rad) and plated at a density of 9.000 cells/well on BIOFLOAT™ 96-well plates (faCellitate, F202003). The plate was centrifuged at 220 × g for 5 min and placed in the incubator at 37 °C/5% CO^2^. After 48 h, the medium was switched to patterning medium (N2B27 supplemented with 200 μM AA, 1 ng/mL BDNF (Prospec, CYT-207), 1 ng/mL GDNF (PeproTech, 450-10), and 1 µM SAG). After two days, the hMLO media was refreshed with patterning medium. After two days, the medium was replaced with maturation medium (N2B27 supplemented with 200 μM AA, 2 ng/mL BDNF, 2 ng/mL GDNF, 1 ng/mL TGF-β3 (Prospec, CYT-368), 5 ng/mL ActivinA (Stem Cell Technologies, 78001), and 100 μM dbcAMP (Sigma-Aldrich, D0260). ActivinA concentration was lowered to 2 ng/mL for the subsequent media changes. The medium was changed every other day. On day 30, the medium was switched to N2B27 without supplements and refreshed every other day.

### Drug treatments

The following reagents were added directly or mixed with growth medium, exosome-free growth medium, or DMEM and placed on cultured cells for the specified time points: Bafilomycin A1 (BafA1, 200 nM, Enzo, BML-CM110-0100), Torin1 (200 nM, Invivogen, inh-tor1), Methylamine hydrochloride (2 mM, Sigma-Aldrich, m0505), Chloroquine (CQ, 50 µM, Sigma-Aldrich, C6628), Brefeldin A (BFA, 2 µg/mL, Cayman Chemical, 11861), Thapsigargin (100 nM, Sigma-Adrich, T9033), Ionomycin (100 nM, Cayman Chemical, 11932), BAPTA-AM (2 µM, Cayman Chemical, 15551), MG-132 (10 µM, Enzo, BML-PI102-0005), GW4869 (4 µM, Sigma-Aldrich, D1692), Dynasore (80 µM, Santa Cruz Biotechnology, sc-202592), Puromycin (Puro, 0.5 - 8 µg/mL Invivogen, ant-pr-1), Doxycycline hyclate (Dox, 1 µg /mL, Sigma-Aldrich, D9891).

### CRISPR/Cas9 LRP10 knock-out in NPCs, HEK-293T, and HuTu-80 cells

To generate LRP10^KO^ NPCs, the Control-1 line was transfected with the pSpCas9-PuroR- LRP10gRNA plasmid using Lipofectamine™ Stem Transfection Reagent (Thermo Fisher Scientific) according to the manufacturer’s specifications. The next day, 0.5 µg/mL of Puromycin (Invivogen) was added to the NPC medium. After two days, scattered single cells were observed and they were allowed to recover with NPC medium without Puromycin. After six days, single colonies were picked and placed in 96-well plates. Recovered clones were expanded as independent clones and genotyped. HEK-293T and HuTu-80 LRP10^KO^ cells were generated according to Grochowska et al. (11, 44). Briefly, HEK-293T and HuTu-80 cells were transfected with the pSpCas9-GFP-LRP10gRNA plasmid using GeneJuice® reagent (Merck) according to manufacturer’s specifications. After two days, cells were dissociated with Trypsin–EDTA (Gibco), and GFP-positive cells were sorted as single cells with the BD FACSAria™ III cell sorter. Recovered clones were expanded as independent clones and genotyped. For the three cell types, we selected biallelically targeted clonal lines carrying a homozygous, single base-pair insertion, predicted to result in a premature stop codon in *LRP10* (p.Gly12Trp fs*18).

### Protein lysates and conditioned media extraction

Cultured cells and hMLOs were washed with PBS and lysed with protein lysis buffer (100 mM NaCl, 1.0% Nonidet P-40 Substitute [Sigma, 74385], 50 mM Tris-Cl, pH 7.4) supplemented with protease inhibitors Complete® (Merck) and Pefabloc® SC (Merck) at 4 °C. For hMLOs, additional mechanical lysis was performed with the Disruptor Genie® (Scientific Industries, Inc.). Lysates were snap-frozen, thawed on ice, and cleared by centrifugation at 10,000 × g for 10 min at 4 °C.

Conditioned media from cultured cells was collected, centrifuged at 10,000 × g for 10 min at 4 °C, and further processed. When specified, cleared conditioned media was concentrated via Amicon Ultra-15 filters (Millipore, UFC901024) and centrifugation for 10-40 min at 4,000 × g at 18 °C. Each filter was used once or twice.

### Western blotting

Protein lysates were mixed with 4 × sample buffer (8% SDS, 20% v/v glycerol, 0.002% bromophenol blue, 62.5 mM Tris-Cl, pH 6.8) supplemented with 400 mM dithiothreitol (DTT) and incubated for 10 min at 95 °C. Conditioned media or EVs fractions were mixed and incubated with 4 × sample buffer without DTT. Proteins were separated on 4-15% Criterion TGX precast gels (Bio-Rad). For experiments including control-3 and patient clone-3 hMLOs, 4-15% Stain-free Criterion TGX precast gels (Bio-Rad) were used, and they were scanned using a Gel Doc XR+ Imaging System (Bio-Rad), to quantify total protein per sample. All gels were transferred to nitrocellulose membranes using the Trans-Blot® Turbo™ Transfer System (Bio-Rad). Blots that were used to detect extracellular α-synuclein were incubated with 0.4% PFA in PBS for 20 min at room temperature (52). All membranes were blocked using 5% non-fat dry milk (Blotto, Santa Cruz Biotechnology) in Tris-buffered saline (TBS), 0.1% v/v TWEEN® 20 (Merck) for 30 min to 1 h at room temperature. Primary antibody incubations were performed in blocking buffer overnight at 4 °C. After washing in TBS, 0.1% v/v TWEEN® 20, blots were incubated for 1 h at room temperature with fluorescently conjugated secondary antibodies. After washing in TBS, 0.1% v/v TWEEN® 20, membranes were imaged and analysed using the Odyssey CLx Imaging system (LI-COR Biosciences).

### Co-immunoprecipitation

After protein lysates were collected, protein concentrations were determined via Pierce™ BCA Protein Assay Kit (Thermo Fisher Scientific). For co-immunoprecipitation, Pierce™ Protein G Magnetic Beads (20 μL, Thermo Fisher Scientific Fisher) were washed three times with washing buffer (50 mM Tris-Cl (pH 7.4), 0.5 M NaCl, 0.05% v/v TWEEN® 20) and incubated with 2 μg of rabbit anti-LRP10 (Sino Biological, 13228-T16) or 5 μg sheep anti-LRP10 (MRC PPU, DA058) antibody for 10 min at RT. Beads were washed three times with washing buffer and incubated with 400 µg protein lysates overnight at 4 °C. For media fractions, 60 µL of EVs or 1 mL of 10 × concentrated supernatant with Amicon filters were used. Beads were washed three times with cold washing buffer and eluted with 30 μL of 1 × sample buffer (2% SDS, 5% v/v glycerol, 0.0005% bromophenol blue, 15.6 mM Tris-Cl (pH 6.8)) supplemented with 100 mM dithiothreitol (DTT) for 10 min at 95 °C. Beads were magnetised and discarded. Samples were analysed by Western blotting.

### Immunocytochemistry of 2D cultures (ICC)

Cells grown on glass coverslips were fixed with 4% PFA for 10 min and washed three times with PBS at room temperature. For uptake experiments in HuTu-80 cells, three additional washings prior to PFA incubation were performed. After washing, cells were incubated in blocking buffer (50 mM Tris.HCl, 0.9% NaCl, 0.25% gelatin, 0.2% Triton™ X-100, pH 7.4) containing primary antibodies at 4 °C overnight. Next, coverslips were washed by dipping them in a PBS, 0.05% TWEEN® 20 solution at room temperature. Cells were then incubated for 1 h with fluorescently conjugated secondary antibodies at room temperature. After washing with PBS, 0.05% TWEEN® 20, cells were mounted with ProLong Gold with DAPI (Invitrogen). For stainings with the cell surface markers AQP4, CD44, and GLAST-1 the blocking buffer consisted of PBS with 2% bovine serum albumin (BSA) and washings were performed with PBS.

### hMLOs immunocytochemistry

hMLOs were collected in tubes, fixed with 4% paraformaldehyde (PFA, Sigma-Aldrich) for 20 min at room temperature, and washed three times with PBS. Next, they were incubated in 30% sucrose in PB buffer (23 mM NaH2PO4(H2O) and 145 mM Na2HPO4(2H2O) in water) at 4 °C overnight. Subsequently, they were embedded in a solution containing 7.5% gelatin and 10% sucrose in PB at room temperature. Solidified gelatin blocks were snap-frozen in cold isopentane and kept at -80 °C. Frozen gelatin blocks were cut into 20 µm sections using a cryostat. For immunocytochemistry, sections were permeabilised and blocked with 0.25% Triton™ X-100 (Sigma-Aldrich), and 4% normal goat serum (Dako) in PBS for 1 h at room temperature. Next, they were incubated with primary antibodies diluted in 0.1% Triton™ X-100, and 4% normal goat serum in PBS at room temperature overnight. The sections were washed three times with PBS and incubated with secondary antibodies diluted in 0.1% Triton™ X-100, and 4% normal goat serum in PBS. After washing three times with PBS, the sections were mounted using ProLong Gold with DAPI (Invitrogen) and imaged on the Leica Stellaris 5 LIA confocal microscope.

### Primary and secondary antibodies

The primary antibodies used for Western Blot (WB) and immunocytochemistry (ICC) are listed in Table 2.

**Table 2:** Primary antibodies used for immunocytochemistry (ICC) and western blotting (WB) experiments. . “*” = This antibody, raised against the N-terminal domain of LRP10, was used for LRP10 detection in all experiments unless otherwise specified.

| Target protein | Host | Supplier | Catalogue number | WB dilution | ICC dilution |
| --- | --- | --- | --- | --- | --- |
| AQP4 | Rabbit | Sigma-Aldrich | HPA014784 | - | 1:100 |
| APC | Mouse | Abcam | AB16794 | - | 1:100 |
| calnexin | Rabbit | Cell Signaling Technologies | 2679S | 1:2000 | - |
| CD44 | Mouse | Abcam | ab6124 | - | 1:100 |
| CD63 | Mouse | ThermoFisher | 10628D | 1:1000 | - |
| CD81 | Mouse | ThermoFisher | 10630D | 1:1000 | - |
| EEA1 | Mouse | BD Biosciences | 610457 | - | 1:200 |
| FOXA2 | Rabbit | Cell Signaling Technologies | 8186 | 1:1000 | - |
| GFAP | Mouse | DSHB | N206A/8 | - | 1:200 |
| GFAP | Rabbit | Sigma-Aldrich | AB5804 | 1:1000 | 1:200 |
| GLAST-1 | Rabbit | Novus | NB100-1869SS | - | 1:100 |
| LRP10* (N-t) | Rabbit | Sino Biological | 13228-T16 | 1:1000 | 1:200 |
| LRP10 (C-t) | Sheep | MRC PPU | DA058 | 1:1000 | - |
| MAP2 | Guinea pig | Synaptic systems | 188 004 | - | 1:500 |
| NANOG | Rabbit | Abcam | AB21624 | - | 1:75 |
| OCT4 | Rabbit | Abcam | AB19857 | - | 1:250 |
| SOX9 | Goat | R&D Systems | AF3075-SP | - | 1:100 |
| SSEA4 | Mouse | Abcam | AB16287 | - | 1:75 |
| s100β | Mouse | Sigma-Aldrich | S2532 | - | 1:200 |
| TGN46 | Sheep | Bio-Rad | AHP500GT | - | 1:300 |
| TH | Sheep | Novus | NB300-110 | 1:1000 | - |
| TRA1-81 | Mouse | Abcam | AB16289 | - | 1:75 |
| tubulin β3 | Mouse | Abcam | AB78078 | 1:5000 | - |
| vinculin | Mouse | Santa Cruz Biotechnology | 2679S | 1:5000 | - |
| α-synuclein | Mouse | BD Biosciences | 610787 | 1:1000 | - |

Secondary antibodies used for Western blotting: Alexa Fluor Plus 680 or 800 donkey anti-mouse and donkey anti-rabbit (all from Thermo Fisher ScientificFisher, 1:1000), or Alexa Fluor® 790 AffiniPure donkey anti-sheep (Jackson immunoresearch, 1:1000).

Secondary antibodies used for immunocytochemistry: Alexa Fluor® Plus 488 donkey anti-rabbit (Thermo Fisher ScientificFisher), Alexa Fluor® 488 anti-mouse/sheep/goat; Alexa Fluor® 594 donkey anti-mouse, Alexa Fluor® 647 goat anti-guinea pig (Jackson ImmunoResearch Laboratories).

Antibodies used for TEM: goat anti-rabbit IgG 15 nm gold-labelled (Aurion, 815.011), ImmunoGold goat anti-mouse 6 nm gold-labelled (Aurion, 806.022).

### Microscopy images acquisition

hMLOs and protein uptake specimens were imaged with the Leica Stellaris 5 LIA confocal microscope. The following lasers were used: diode 405, OPSL 488, DPSS 561, and diode 638. Each image was detected with the 3 × spectral HyD S detector with an HC PL APO CS2 40 ×/1.3 or HC PL APO CS2 63 ×/1.4 lens. The rest of the specimens were imaged with Leica SP5 AOBS confocal microscope. The following lasers were used: diode 405, OPSL 488, DPSS 561, and HeNe 633. Each image was detected on the spectral PMT detector with an HCX PL APO CS 40 ×/1.25 or HCX PL APO CS 63 ×/1.4 lens. For both microscopes, scanning of detailed images was done with a pixel size of 0.1 μm and with a scan size of 2048 × 2048 pixels at 400 or 600 Hz. For z-stack images, 0.35 μm steps in the z-direction were taken. The pinhole size was set to 1 airy unit (AU). Scanning of overview hMLOs images consisted of a tile scan of 9 regions with a pixel size of 0.285 μm, a scan size of 1024 × 1024 pixels at 600 Hz, and 0.5 μm steps in the z-direction.

### Image Analysis

Microscopy images were analysed using Fiji/ImageJ version 1.45b. For colocalisation of LRP10 and cellular markers in control hMLOs (Fig. 3A-E), big artifacts and background signal were removed from each channel using automatic thresholding and size filtering. The built-in tool coloc2 was used to determine the Manders’ coefficient. The channels overlap of each Z plane was averaged for each hMLO. The rest of analyses were performed on maximum projections. To quantify LRP10 uptake in HuTu cells (Fig. 2), a mask for cell surfaces was generated using the LRP10 antibody channel including background signal from the cell cytoplasm. LRP10 signal was then measured from individual cells after automatic thresholding and generating a mask exclusively containing overlapping signal from the LRP10 antibody and mCherry channels. To quantify LRP10 in neurons in DMSO and BafA1-treated hMLOs (Fig. 3F and 3G), 30 MAP2- positive neurons were randomly selected from two hMLOs per condition, and their cell surface was manually outlined after generating short maximum projections (3-5 planes of 0.5µm each) that were adjusted per cell. LRP10 signal intensity was measured from each cell. To study LRP10 levels in s100β-positive cells in control and patient hMLOs (Fig. 6C and 6D), a mask for s100β was generated after automatic thresholding. LRP10 objects within the s100β mask were measured after an additional automatic thresholding step. To quantify the mean intensity and area % of LRP10 wild-type (LRP10^WT^) and the patient-derived LRP10 splice variant (LRP10^splice^) in overexpressing cells (Fig. 7A-C), the cell surfaces of single cells were selected using the LRP10 antibody channel including background signal from the cell cytoplasm, and LRP10 signal was quantified per cell after automatic thresholding.

**Figure 1:**
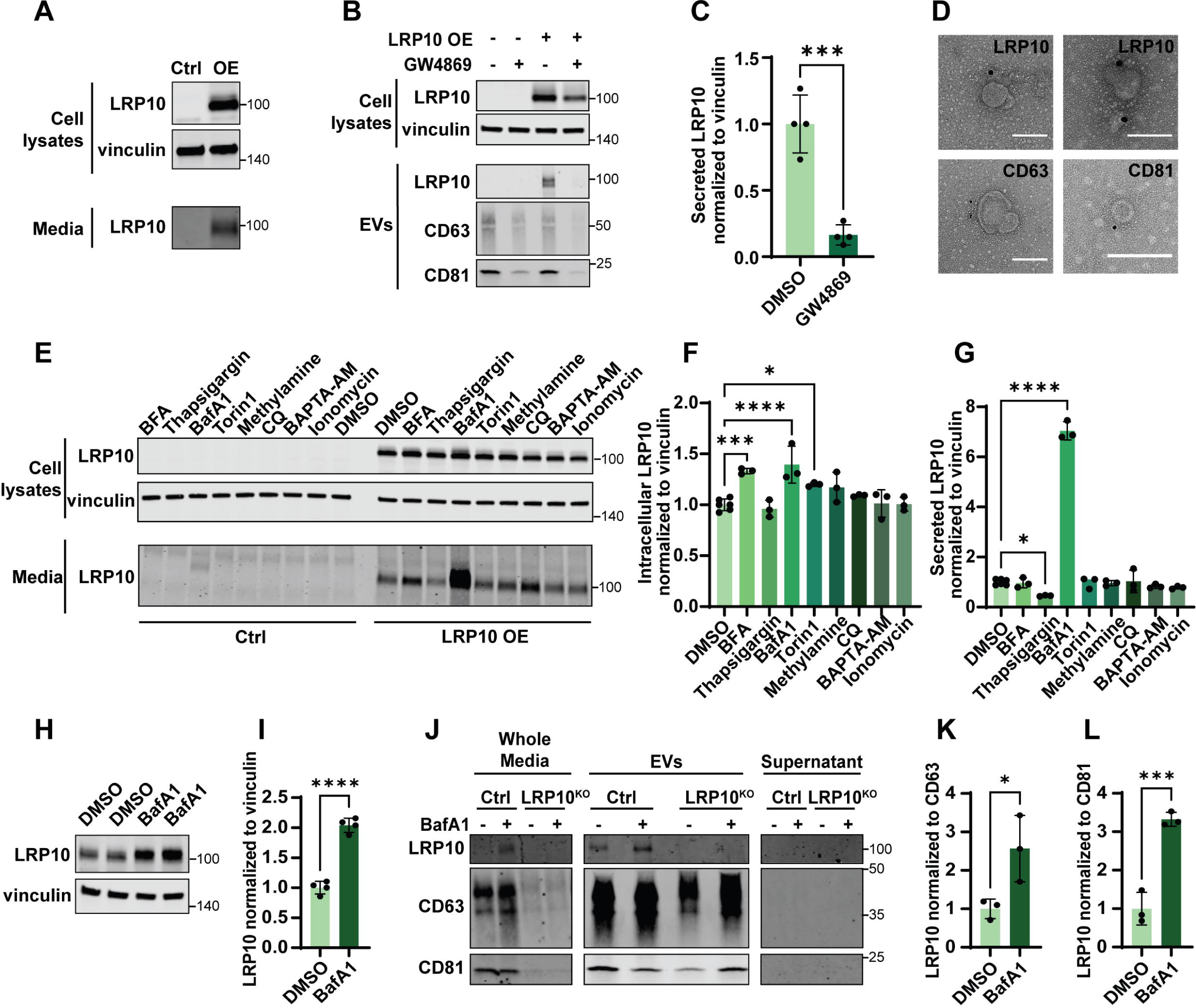
LRP10 secretion mechanisms. (**A**) Representative western blot of cell lysates and conditioned media from control (Ctrl) and LRP10-overexpressing (OE) HEK-293T cells probed for LRP10 and vinculin. (**B**) Representative western blot of cell lysates and EVs isolated from conditioned media from LRP10-overexpressing cells after 48 h of DMSO or GW4869 treatments. Blots were probed for expression of LRP10, vinculin, CD63, and CD81. (**C**) Western blot quantification of LRP10 in EVs normalised to vinculin in cell lysates from (**B**). N = 4 biological replicates. (**D**) Representative TEM images of LRP10, CD63, and CD81 immunoreactive EVs. Scale bars = 100 nm. (**E**) Representative western blot of LRP10-overexpressing cells treated with the indicated compounds for 4 h. (**F**, **G**) Western blot quantifications of intracellular (**F**) and extracellular (**G**) LRP10 normalised to intracellular vinculin from (**E**). N = 3 biological replicates. (**H**, **I**) Representative western blot (**H**) and quantification (**I**) of endogenous LRP10 in 2 months- old control iPSC-derived astrocytes treated with DMSO or BafA1 for 4 h. N = 4 biological replicates. (**J**) Representative western blot of fractionated conditioned media from 4 months-old control and LRP10^KO^ iPSC-derived astrocytes after 2 days in culture with exosome-free media followed by a 6 h treatment with DMSO or BafA1. Conditioned media (Whole Media) was fractionated into EVs and supernatant fractions. (**K**, **L**) Quantification of LRP10 in EVs normalised to the EVs markers CD63 (**K**) and CD81 (**L**) from (**J**). N = 3 biological replicates. All data are expressed as mean ± SD with individual data points shown. Data in (**C**, **I**, **K**, **L**) were analysed by unpaired T-test. Data in (**F**, **G**) were analysed by One-way ANOVA with Dunnett’s multiple comparisons test. *\*P* ≤ 0.05, \*\*\**P* ≤ 0.001, \*\*\*\**P* ≤ 0.0001.

**Figure 2:**
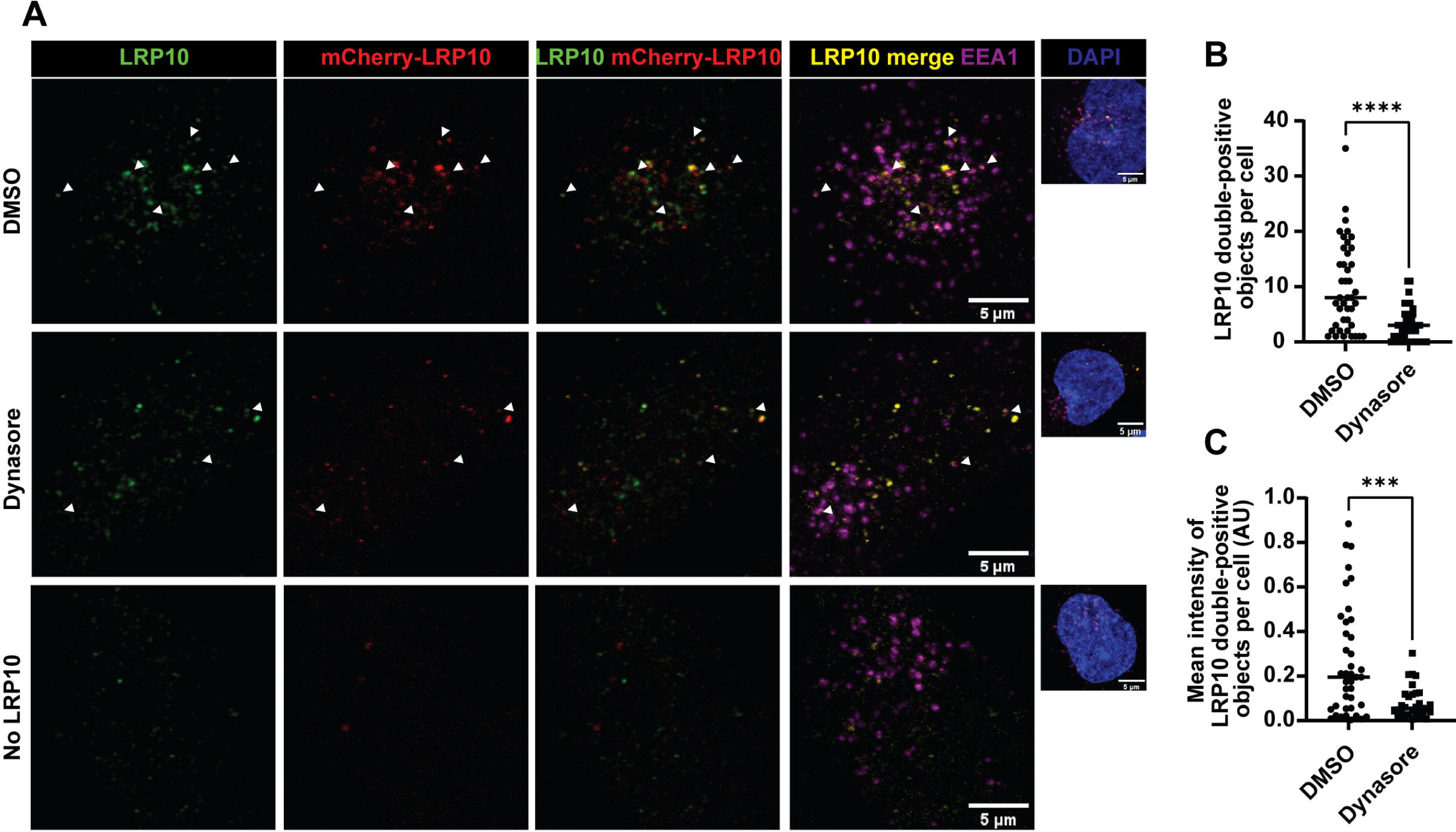
LRP10 uptake via clathrin-mediated endocytosis. (**A**) Representative confocal images of LRP10 and mCherry-LRP10 internalised by DMSO or Dynasore-treated LRP10^KO^ HuTu-80 cells. White arrowheads indicate LRP10 in EEA1-positive vesicles. (**B**, **C**) Quantifications of LRP10 double-positive objects (**B**) and their mean signal intensity (**C**) per cell. N ≥ 40 biological replicates. All data are expressed as mean ± SD with individual data points shown. Data were analysed by unpaired T-test. \*\*\**P* ≤ 0.001, \*\*\*\**P* ≤ 0.0001.

**Figure 3:**
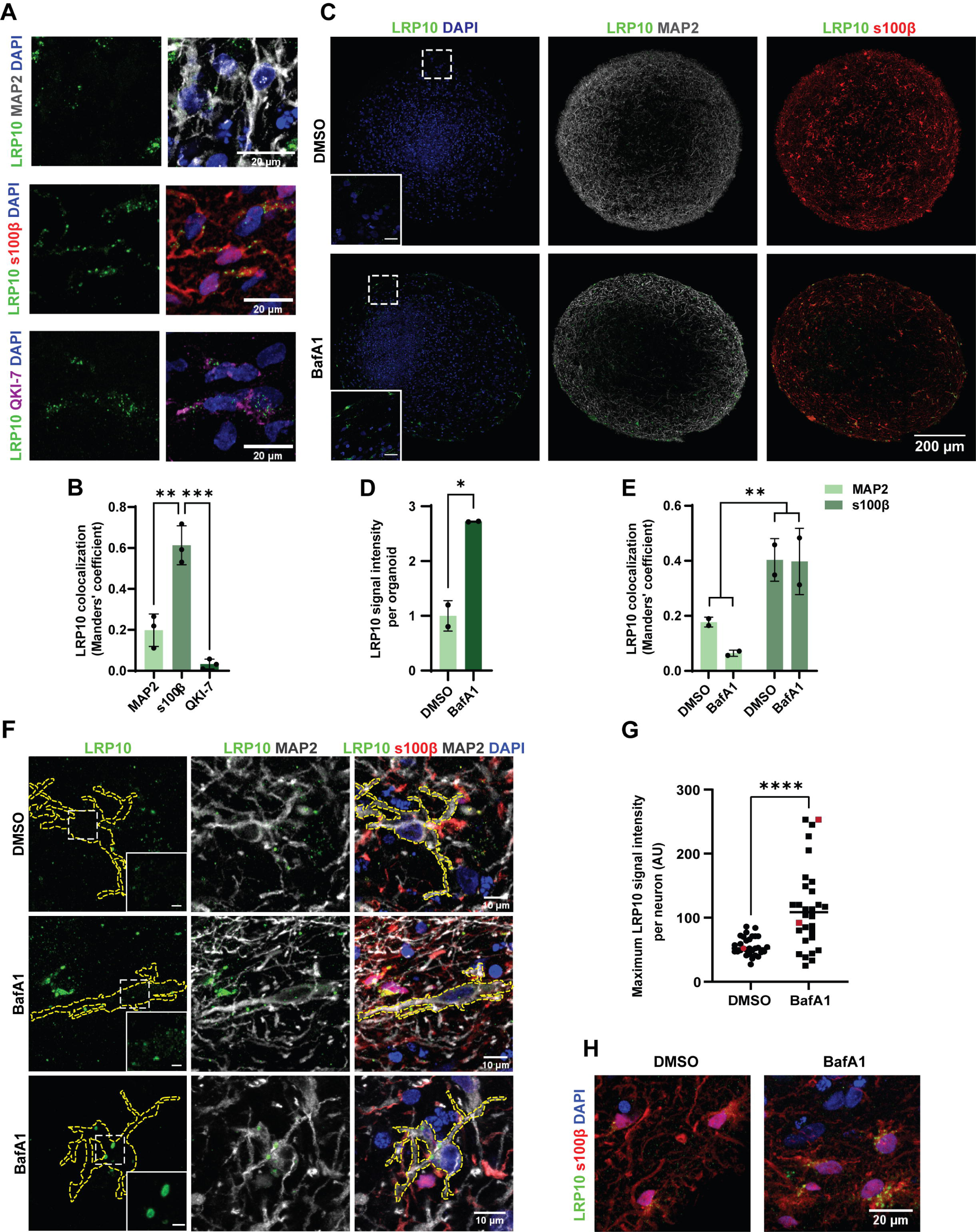
LRP10 localisation in untreated and BafA1-treated hMLOs. (**A**) Representative confocal images of LRP10 localisation in 2 months-old control, untreated hMLOs relative to the cellular markers MAP2 (neurons), s100β (astrocytes), and QKI-7 (oligodendrocytes). (**B**) Colocalisation analysis via Manders’ overlap of LRP10 with MAP2, s100β, and QKI-7 from (**A**). N = 3 biological replicates. (**C**) Representative tile scans of LRP10 in 4 months-old control hMLOs treated with DMSO or BafA1 for 4 h relative to the markers MAP2 and s100β. White dotted boxes indicate the zoomed-in regions in the bottom left corners (Scale bars = 20 µm). (**D**, **E**) Quantification of LRP10 signal intensity (**D**) and LRP10 colocalisation with MAP2 (neurons) and s100β (astrocytes) (**E**) per hMLO from (**C**). N = 2 biological replicates. (**F**) Representative detailed confocal images of LRP10 in MAP2-positive neurons in DMSO or BafA1-treated hMLOs from (**C**). The cell surface of single MAP2-positive cells is indicated with yellow dotted lines. White dotted boxes indicate the zoomed-in regions for LRP10 in the bottom right corners for the LRP10 channel (Scale bars = 2 µm). (**G**) Quantification of LRP10 maximum signal intensity per MAP2- positive neuron. Red data points indicate the values of the representative cells shown in (**F**). N = 30 biological replicates. (**H**) Representative detailed confocal images of LRP10 in s100β-positive astrocytes in DMSO or BafA1-treated hMLOs from (**C**). All data are expressed as mean ± SD with individual data points shown. Data from (**B**) were analysed by One-way ANOVA with Tukey’s multiple comparisons test. Data from (**D**, **G**) were analysed by unpaired T-test. Data from (**E**) were analysed by Two-way ANOVA. *\*P* ≤ 0.05, \*\**P* ≤ 0.01, \*\*\**P* ≤ 0.001, \*\*\*\**P* ≤ 0.0001.

### EVs isolation from conditioned media

HEK-293T cells were grown in T75 cell culture flasks and refreshed two days before media collection with exosome-free growth media (FBS was replaced by One Shot™ Fetal Bovine Serum, Exosome-Depleted, Gibco) supplemented with GW4869 or DMSO when indicated. Ten millilitres of conditioned media were collected and centrifuged at 2,000 × g for 10 min at 18 °C to remove cell debris. The initial supernatant (also referred to as “whole media” in Fig. 1J and Supplementary Fig. 1) was collected and further centrifuged at 16,000 × g for 20 min at 4 °C to remove big vesicles. The resulting supernatant was filtered with a membrane pore size of 0.2 μm (Whatman) and centrifuged at 120,000 × g for 4 h at 4 °C in polycarbonate bottle assemblies (Beckman coulter, 355603) in a 70.1 Ti fixed angle rotor (Beckman coulter, 342184) in an Optima XE-90 ultracentrifuge (Beckman coulter) to collect smaller EVs, including exosomes. The final supernatant was collected as the “supernatant” fraction and the pellet was resuspended with 70 μL of PBS (“EVs” fraction). EVs were further processed or stored at -80 °C.

iPSC-derived astrocytes were grown in T25 flasks with exosome-free N2 medium (FBS was replaced by One Shot™ Fetal Bovine Serum, Exosome-Depleted, Gibco), and conditioned media was collected and frozen at -80 °C four consecutive times with a two-day interval. Twenty millilitres of media were then thawed on ice, centrifuged at 2,000 × g for 10 min at 18 °C, and the supernatant was concentrated until reaching a volume of 10 mL with Amicon filters. The concentrate was collected and further centrifuged at 16,000 × g for 20 min at 4 °C, filtered with a 0.2 μm membrane, and ultracentrifuged at 120,000 × g for 4 h at 4 °C. The final supernatant was collected (“supernatant” fraction) and the pellet (“EVs fraction”) was resuspended with 70 μL of PBS.

### EV quantification (EVQuant)

EVs were quantified using the recently developed EVQuant assay, described previously (53). Briefly, isolated EVs, supernatant, and whole media fractions were mixed with the fluorescent membrane dye Octadecyl Rhodamine B Chloride R18 (0.33 ng/ul, Life Technologies, O246) and incubated for 10 min at room temperature. Next, fluorescently labelled EVs were immobilised in a non-denaturing polyacrylamide gel (16% w/w acrylamide/bis-acrylamide) in 96-wells glass-bottomed plates (Cellvis, P96-0-N). Per sample, 25 single-plane images were acquired using a high-content screening system (Opera Phenix, Perkin Elmer) using a 40x water immersion objective (NA 1.1). Rhodamine-labelled nanoparticles were excited using a 561nm laser and detected at 570-630nm emission wavelength. EV concentration was calculated as the number of detected EVs in a calibrated confocal imaging volume (53). EV concentrations were background corrected using a dye-only control (sample containing PBS mixed with Rhodamine).

### Transmission electron microscopy (TEM)

EVs purified from conditioned media were evaluated by TEM. To prepare samples for TEM, 10 µl of isolated EVs were incubated for 20 min with a formvar/carbon-coated 400 mesh copper grid. Next, the grids were washed 3 times with PBS, incubated for 30 min with 10 µl of blocking solution containing normal goat serum (Aurion, 905.002), and washed two times with 0.1% BSA- c (Aurion, 900.099). Primary antibodies (rabbit anti-LRP10, mouse anti-CD81, or mouse anti- CD63, 1:50) were diluted in 0.1% BSA-c and added to the samples for 1 h. The grids were washed four times with 0.1% BSA-c and incubated for 1 h with the gold-coupled secondary antibodies diluted in 0.1% BSA-c (1:50 dilution). Next, they were washed five times with 0.1% BSA-c, three times with PBS, and three times with ultrapure water. Grid staining was performed with Uranyless EM Stain for 1 min (negative stain). Grids were air-dried and visualised with a TALOS L120C TEM at 120 kV at 11 k–45 k magnification.

### Nanoparticle tracking analysis (NTA)

NTA was used to quantify and characterise isolated EVs from iPSC-derived astrocytes or exosome-free astrocyte media (Advanced DMEM/F-12, 4% exosome-free FBS, 1% N2, 1% Penicillin-Streptomycin), and HEK-293T cells or exosome-free HEK-293T media (DMEM, 10% exosome-free FBS, 1% Penicillin-Streptomycin) with the NanoSight NS3000 system (Malvern, Sysmex) in light scatter mode using the 405 nm laser. Samples were diluted 1:10 in DPBS (Gibco) and three consecutive measurements were performed for 60 seconds at a controlled temperature of 25°C.

### Conditioned media uptake

HEK-293 cells were cultured in growth medium in 10 cm cell culture dishes. The day before media collection, cells were refreshed with DMEM. Next, 10 mL of conditioned media was collected and centrifuged at 2,000 × g for 10 min. The supernatant was concentrated 100 × with Amicon concentrating filters and mixed with fresh media until reaching a final concentration of 10 ×. For uptake of LRP10 in HuTu-80 cells, cells were washed with PBS and refreshed with DMEM:F12 (Gibco, 11320-074) supplemented with DMSO or 80 µM Dynasore for 30 min, prior to the addition of 1:10 conditioned media concentrate for 30 additional min before further processing. For additional uptake experiments, 1:10 conditioned media concentrate was mixed with growth medium and placed on the donor cells for the specified time points.

### Statistical analysis

Statistical analyses were performed using Prism 9 software (GraphPad). For experiments with only two conditions, students T-test or Mann Whitney test were used. For experiments with two or more groups One-way or Two-way ANOVA followed by the Dunnett’s, Tukey’s, or Sidak’s multiple comparisons tests were used. The Grubb’s test was used to discard outliers. Significant P values of *P* ≤ 0.05 were reported. In the graphs, “*” represents a p-value of *P* ≤ 0.05, “**” represents *P* ≤ 0.01, “***” represents *P* ≤ 0.001, and “****” represents *P* ≤ 0.0001. The data are presented as means ± standard deviations (SD) and represent results from at least 2 biological replicates.

## Results

### LRP10 is secreted via extracellular vesicles and is sensitive to autophagic function

Work from Grochowska et al. (11) revealed that LRP10 is mainly expressed in non-neuronal cells in adult human brain. However, in LBD cases, LRP10 was also detected in Lewy bodies within dopaminergic neurons. To address whether LRP10 in neuronal Lewy bodies possibly originates from non-neuronal cells, we first investigated LRP10 secretion in cultured cells. We studied LRP10 distribution in cell lysates and conditioned media from LRP10-overexpressing (OE) HEK-293T cells by western blot analyses. As a result, LRP10 was detected in both fractions 48 h after transfection (Fig. 1A). LRP10 is a transmembrane protein, so we hypothesised that LRP10 would partition to a lipid membrane-containing fraction, potentially EVs. To address this issue, we fractionated conditioned media (referred to as “whole media”) via sequential centrifugation, filtration, and ultracentrifugation, which resulted in a pellet -consisting of small EVs (54)- and supernatant fractions (Supplementary Fig. 1A). Effectiveness of the isolation was demonstrated via different methods (54). First, western blot analyses revealed the presence of the EVs markers CD63 and CD81 and the absence of the ER-resident protein calnexin in the whole media and EVs fractions (Supplementary Fig. 1B). The absence of calnexin in media fractions also indicated the lack of cell death-related contamination. Next, particle count quantification via membrane-labelling via EVQuant (53) demonstrated the successful concentration of EVs from whole media (Supplementary Fig. 1C; LRP10 wild-type (LRP10^WT^) whole media mean: 4 × 10^10^ particles; supernatant mean: 9 × 10^9^ particles; EVs mean: 4 × 10^11^ particles). Finally, nanoparticle tracking analysis (NTA) was used to determine the size of the purified EVs, which presented a mean diameter of 135.5 nm ± 69.7 and a mode of 84.5 nm (Supplementary Fig. 1D), which is in line with the expected size of exosomes (50-150 nm) and other EVs subtypes (54). Furthermore, NTA also showed an enrichment of vesicles after EVs purification from conditioned media in comparison to media that had not been in contact with cells (Supplementary Fig. 1D). To prove that LRP10 is secreted via EVs, we used LRP10-overexpressing cells and treated them with DMSO or GW4869, a compound that inhibits exosome release. After two days, protein extracts and conditioned media were collected, and EVs were isolated. LRP10 was found in the EVs fraction (Fig. 1B and Supplementary Fig. 1B) but not in the supernatant fraction (Supplementary Fig. 1B). Furthermore, LRP10 levels in EVs were significantly lower after GW4869 treatment with a 6-fold reduction (Fig. 1C). To further confirm the presence of LRP10 in EVs, we performed immuno-transmission electron microscopy (TEM) and labelled LRP10, CD63, and CD81 with golden particles in isolated EVs from conditioned media (Fig. 1D). This assay also confirmed the successful isolation of vesicular structures of the expected sizes (diameter of 50-150 nm for exosomes) (54). Taking these observations together, we conclude that LRP10 is secreted via EVs.

To gain more mechanistic insight into LRP10 secretion, we treated control and LRP10-overexpressing HEK-293T cells for 4 h with compounds that alter cellular functions involved in different secretion pathways and studied their effect on the intra- and extracellular distribution of LRP10 via western blot (Fig. 1E). Brefeldin A (BFA), an ER-to-Golgi vesicle transport and conventional secretion inhibitor, induced a significant increase in intracellular levels of LRP10, but did not affect extracellular LRP10 levels, suggesting that LRP10 follows an unconventional secretory pathway (Fig. 1E-G). Thapsigargin, an ER stressor that blocks Ca^2+^ entry into the ER, did not change the amount of intracellular LRP10 but decreased LRP10 extracellular levels. BafA1, a vacuolar H^+^ ATPase inhibitor that blocks vesicle acidification and autophagy, increased LRP10 intracellular levels and induced a striking 7-fold increase in LRP10 secretion (Fig. 1E-G).

Additionally, BafA1 treatment also led to the detection of endogenous LRP10 in conditioned media from control cells. Torin1, which promotes autophagic function, led to a decrease in intracellular LRP10 without affecting extracellular LRP10 levels. Methylamine and chloroquine (CQ), which inhibit vesicle acidification by different mechanisms, had no effect on intra- and extracellular LRP10 levels, nor did the Ca^2+^ level modifiers BAPTA-AM and Ionomycin (Fig. 1E-G). Taken together, these results show that LRP10 is secreted via EVs, and can be significantly enhanced after blockade of the autophagy pathway.

Due to their relatively high LRP10 expression levels (11, 55–57), we used a previously established *in vitro* iPSC-derived astrocytes model (11, 49) to study the effect of BafA1 treatment on LRP10 under more physiological conditions. Successful generation of control astrocytes derived from a non-demented donor and knock-out (LRP10^KO^) astrocytes was confirmed via immunocytochemistry for the astrocytic markers glial fibrillary acidic protein (GFAP), S100 calcium-binding protein β (s100β), SRY-box transcription factor 9 (SOX9), aquaporin 4 (AQP4), CD44, and Glutamate/Aspartate transporter 1 (GLAST-1) (Supplementary Fig. 2). To determine whether autophagic inhibition affects endogenous LRP10 intracellular levels, control astrocytes were treated with BafA1 for 4 h, and cell lysates were analysed via western blotting (Fig. 1H). As a result, we observed a 2-fold increase of LRP10 intracellular levels after BafA1 treatment (Fig. 1I). Next, to study LRP10 secretion, control and LRP10^KO^ astrocytes were cultured for two days in exosome-free media and treated with DMSO or BafA1 6 h before conditioned media was collected and fractionated. Western blot analyses revealed the presence of LRP10 in EVs from control astrocytes, but not in the supernatant fraction (Fig. 1J). Furthermore, we observed a significant 2.6 to 3.3-fold increase in LRP10 in EVs in BafA1-treated control astrocytes (Fig. 1K and 1L). Strikingly, LRP10 was already detected in whole media from BafA1-treated control astrocytes (Fig. 1J). In addition to presenting CD63 and CD81 (Fig. 1J), the purified EVs were also characterised via NTA (Supplementary Fig. 1E), which showed a peak in the EVs diameter of 140.5 nm. To conclude, autophagic inhibition induces an increase in LRP10 intracellular levels and secretion, particularly via EVs, in iPSC-derived astrocytes, which is in line with our observations using LRP10-overexpressing cells (Fig. 1E-G).

### LRP10 is internalised via clathrin-mediated endocytosis

We next investigated whether secreted LRP10 can be internalized by acceptor cells. We used mCherry-LRP10-overexpressing HEK-293T cells to produce and secrete mCherry-LRP10 into conditioned media, and after 48 h we placed it onto Dynasore (clathrin-mediated endocytosis inhibitor) (58)-treated LRP10^KO^ cells for 30 min. Recipient cells were subsequently inspected via immunocytochemistry for the presence of double-positive vesicles containing both mCherry signal and signal from our KO-validated LRP10 antibody to increase specificity (Fig. 2A) (11). We observed several double-positive vesicles that were also colocalising with the endosomal marker early endosome antigen 1 (EEA1) (Fig. 2A). Additionally, a significantly lower number of double-positive vesicles were detected after Dynasore treatment (Fig. 2B), which also presented lower LRP10 signal intensity (Fig. 2C). These findings suggest that LRP10 can be internalised via clathrin-mediated endocytosis.

### LRP10 expression is enriched in astrocytes in hMLOs

To address whether the origin of LRP10 in Lewy bodies could be the result of an interaction between different brain cell types, we turned to hMLOs as a model system. Control iPSCs were differentiated into hMLOs according to published protocols (50, 51). Even though these hMLOs are described to be partially functional after 1 month *in vitro*, astrocyte markers start appearing after 2 months in culture (51). This observation is in line with normal neurodevelopment, where astrocytes are specified later than neurons (59). Therefore, we characterised our hMLOs at 2 and 4 months after the start of differentiation by immunocytochemistry and western blotting analyses (Supplementary Fig. 3). We detected various cell types, including neurons (Supplementary Fig. 3A), astrocytes (Supplementary Fig. 3B), and oligodendrocytes (Supplementary Fig. 3C), in addition to the presence of tyrosine hydroxylase (TH) and forkhead box protein A2 (FOXA2), a developmental midbrain-specific marker (60) (Supplementary Fig. 3D and 3E). From 2 to 4 months in culture we observed increased levels of several markers, including LRP10 (1.7-fold, not significant), FOXA2 (4-fold, not significant), and the astrocytic marker GFAP (18-fold, \*\**P* = 0.0026). Therefore, 4 months-old hMLOs were selected for further testing due to their higher astroglial levels.

To investigate LRP10 cellular localisation and test whether the findings on LRP10 restricted cell-type expression pattern in human brain material could be reproduced in hMLOs, we performed immunocytochemistry and subsequent colocalisation analysis between LRP10 and different cellular markers in control hMLOs (Fig. 3A and 3B). As a result, LRP10 showed a 20% overlap with the neuronal marker microtubule-associated protein 2 (MAP2), a 61% overlap with the astrocytic marker s100β, and a 3% overlap with the oligodendrocytic marker Quaking 7 (QKI-7) (61) (Fig. 3B). Therefore, LRP10 localisation in hMLOs is enriched in astrocytes, which is in line with previous findings in human post-mortem brain material (11).

### Autophagy inhibition induces the formation of atypical LRP10 objects in neurons of hMLOs

Accumulating evidence suggests that defective autophagic function might play an important role in PD and DLB pathogenesis (62). Since we show that inhibition of the autophagy pathway via BafA1 treatment leads to increased LRP10 secretion, we speculate that defective autophagy in LBD brains could be a potential cause for enhanced LRP10 secretion from non-neuronal cells and subsequent neuronal uptake and accumulation in Lewy bodies. To investigate this possibility, we treated control hMLOs with DMSO or BafA1 for 4 h and studied LRP10 levels and localisation via immunocytochemistry (Fig. 3C-H). First, we observed 1.7-fold higher LRP10 levels in whole hMLOs after BafA1 treatment (Fig. 3C and 3D), which is in line with the observed effect of BafA1 on LRP10-overexpressing cells and astrocytes (Fig. 1E-I). Inspection of LRP10 localisation relative to the astrocytic marker s100β and the neuronal marker MAP2, revealed that LRP10 remained enriched in astrocytes in comparison to neurons before and after BafA1 treatment (Fig. 3C, 3E, and 3H). However, analysis of single MAP2-positive neurons showed a significant 2-fold increase in LRP10 signal after BafA1 treatment, as well as neurons with atypical enlarged LRP10+ intracellular vesicular structures (Fig. 3F and 3G). These data show that autophagic stress induces an increase in LRP10 levels in hMLOs in astrocytes and neurons, and the formation of atypical LRP10 vesicular structures in neurons.

### LRP10 regulates α-synuclein intracellular levels and secretion

The aggregation and associated neuronal toxicity of α-synuclein are considered to be central in PD and DLB pathogenesis. Since both LRP10 and α-synuclein are found in Lewy bodies (11), we sought to study a possible functional link between LRP10 and α-synuclein. First, we overexpressed variable amounts of both proteins (encoded by *LRP10* and *SNCA*, respectively) in HEK-293T cells to investigate their interaction after two days via western blot (Fig. 4A). Interestingly, α-synuclein overexpression led to a significant decrease in intracellular LRP10 levels (Fig. 4B), and, in contrast, LRP10 overexpression induced increased intracellular α-synuclein levels (Fig. 4C). The strongest effect was found in the media fraction, where we observed a striking 150 to 300-fold increase in extracellular α-synuclein levels in the condition of combined α-synuclein and LRP10 overexpression (Fig. 4D). These results indicate that LRP10 and α-synuclein have an opposite, reciprocal effect on the intracellular levels of the other protein, and that LRP10 promotes α-synuclein secretion.

**Figure 4:**
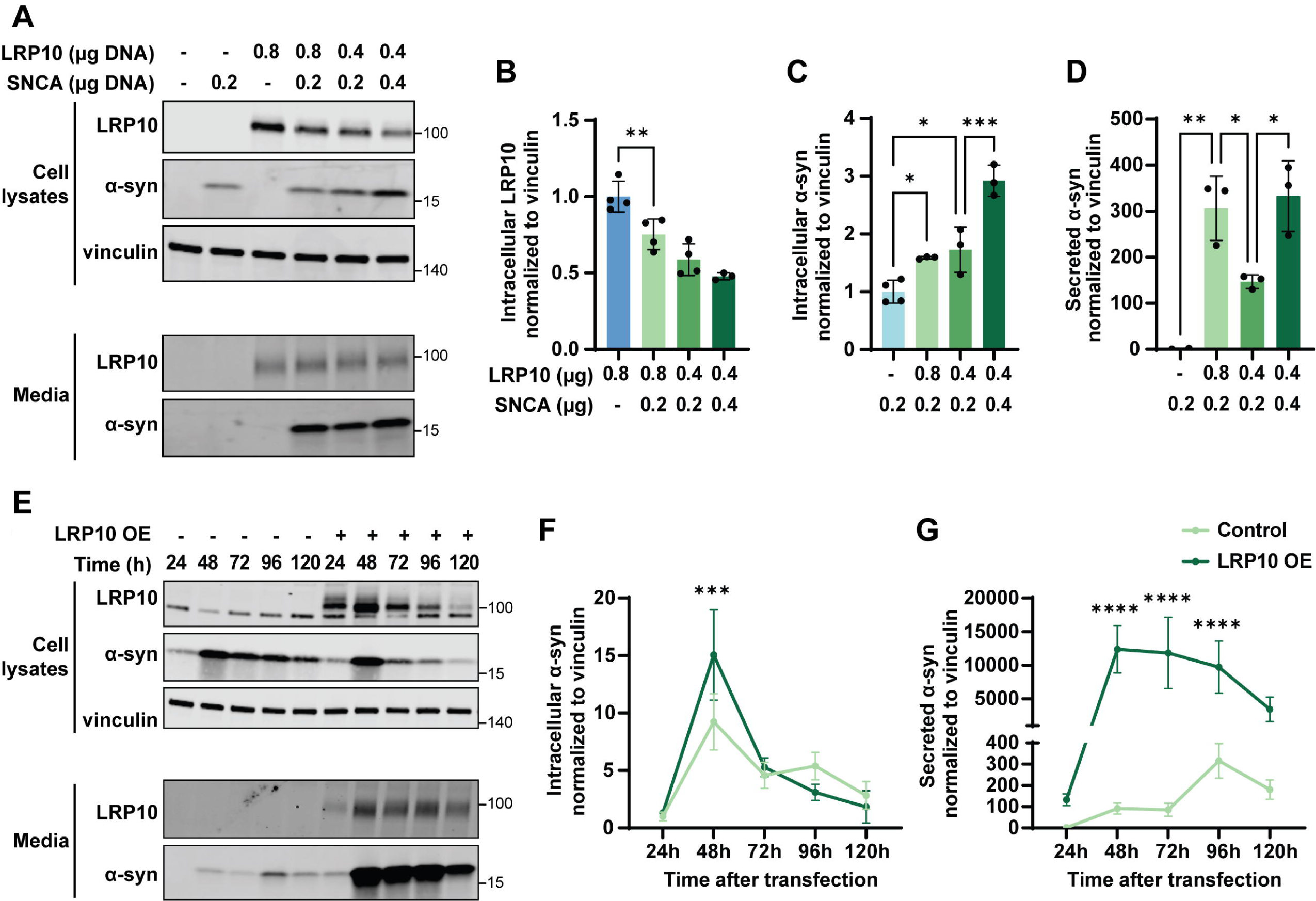
LRP10 and α-synuclein interaction. (**A**) Representative western blot of cell lysates and conditioned media from HEK-293T cells transfected for 48 h with different amounts of *LRP10* and *SNCA.* (**B**-**D**) Western blot quantifications of intracellular LRP10 (**B**), intracellular α-synuclein (α-syn) (**C**), and secreted α-synuclein normalised to intracellular vinculin (**D**). N = 4 biological replicates. (**E**) Representative western blot of cell lysates and conditioned media from LRP10 and α-synuclein-overexpressing cells over time. (**F**, **G**) Western blot quantifications of intracellular α-synuclein (**F**) and secreted α-synuclein (**G**) normalised to intracellular vinculin. N = 3 biological replicates. All data are expressed as mean ± SD with individual data points shown in (**B**-**D**). Data in (**B**-**D**, **F**, **G**) were analysed by One-way ANOVA with Sidak’s multiple comparisons test. *\*P* ≤ 0.05, \*\**P* ≤ 0.01, \*\*\**P* ≤ 0.001, \*\*\*\**P* ≤ 0.0001.

To better understand the dynamics of the LRP10-α-synuclein interaction, we examined the effect of LRP10 overexpression on intra- and extracellular α-synuclein levels over time using the same experimental paradigm (Fig. 4E). LRP10 overexpression led to elevated intracellular α-synuclein levels at 48 h post-transfection when compared to control cells, but this initial effect was reversed from 72 h onwards (Fig. 4F). In the media fraction, control cells showed a slight increase in α- synuclein levels from 96 h onwards, whereas LRP10-overexpressing cells showed a peak in extracellular α-synuclein levels at 48 h post-transfection (Fig. 4G). Nonetheless, LRP10-overexpressing cells showed higher extracellular α-synuclein levels than control cells overall (LRP10 OE vs Control: \*\*\*\**P* < 0,0001). To conclude, LRP10 dynamically regulates intracellular α-synuclein levels over time and strongly promotes its secretion.

### LRP10 induces α-synuclein secretion via unconventional secretion pathways

Several studies have shown that α-synuclein is secreted via a variety of pathways, including EVs (63, 64). To determine whether LRP10 promotes α-synuclein secretion via EVs, we purified EVs from conditioned media of LRP10 and α-synuclein-overexpressing cells treated with DMSO or GW4869 for 48 h. Western blotting revealed the presence of α-synuclein in the EVs and the supernatant fractions (Fig. 5A). In both fractions, LRP10 overexpression induced a significant increase in α-synuclein levels (Fig. 5B). In contrast to the inhibitory effect of GW4869 treatment on LRP10 localisation in EVs, GW4869 induced an increase in α-synuclein levels in EVs, which suggests the activation of an alternative secretion pathway (Fig. 5A and 5B, DMSO vs GW4869 \**P* = 0.0305). Additionally, GW4869 treatment did not affect α-synuclein secretion differently in LRP10-overexpressing cells in comparison to control cells (Fig. 5C). These data indicate that LRP10 induces α-synuclein secretion via EVs and EVs-independent mechanisms.

**Figure 5:**
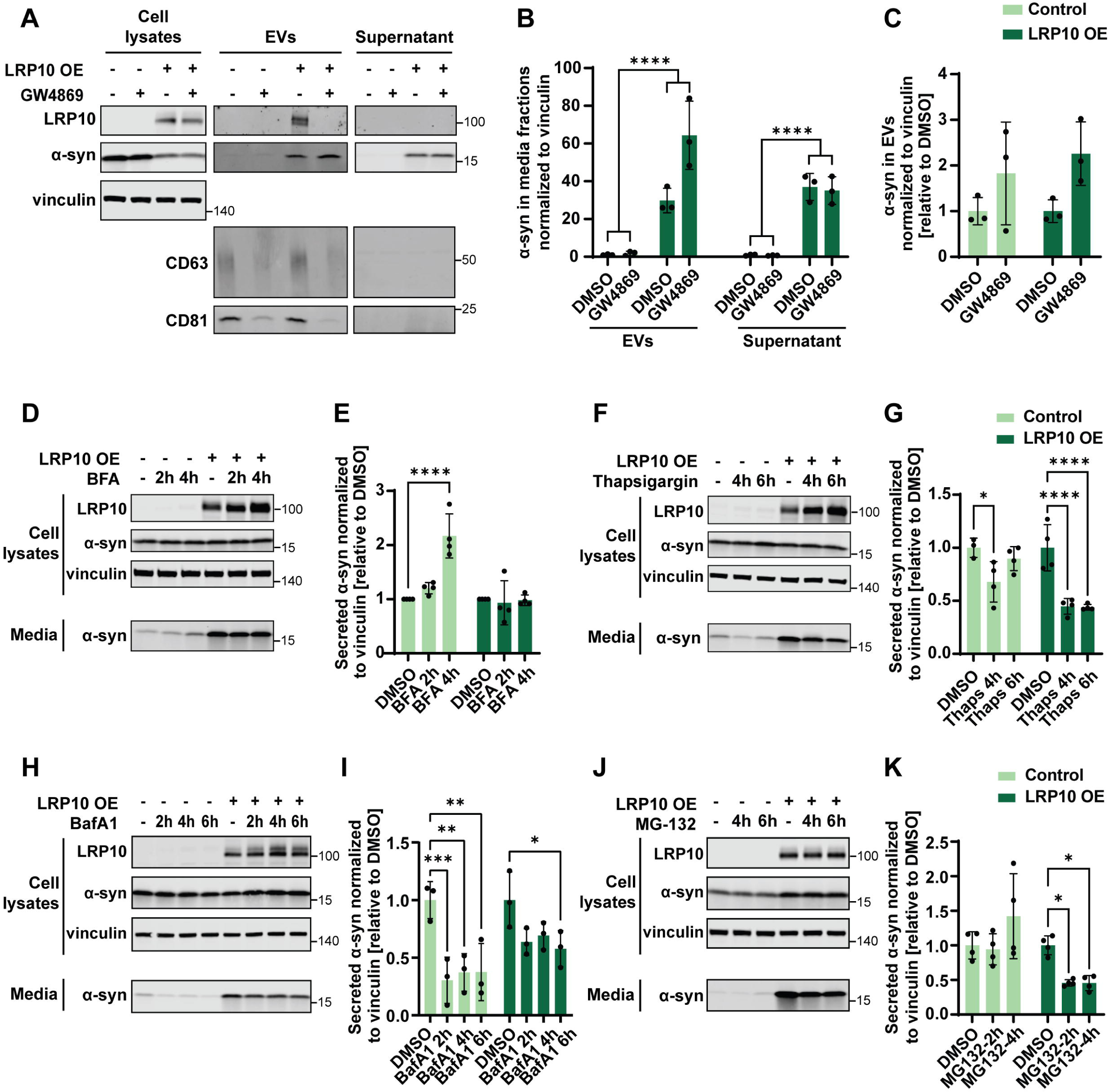
LRP10-mediated α-synuclein secretion mechanisms. (**A**) Representative western blot of cell lysates and media fractions (EVs and Supernatant) from 48 h DMSO or GW4869-treated LRP10 and α-synuclein-overexpressing cells. (**B**, **C**) Western blot quantifications of α-synuclein from EVs and Supernatant fractions normalised to vinculin (**B**) and α-synuclein from the EVs fraction normalised to vinculin relative to DMSO-treated conditions (**C**). N = 3 biological replicates. (**D**-**K**) Representative western blots and quantifications of LRP10 and α-synuclein-overexpressing cells treated with BFA (4 h secretion in total; **D**, **E**), Thapsigargin (6 h secretion in total; **F**, **G**), BafA1 (6 h secretion in total; **H**, **I**), and MG-132 (6 h secretion in total; **J**, **K**). N = 3 or 4 biological replicates. All data are expressed as mean ± SD with individual data points shown. Data in (**B**, **C**) were analysed by Two-way ANOVA. Data in (**E**, **G**, **K**) were analysed by Two-way ANOVA with Sidak’s multiple comparisons test. Data in (**I**) were analysed by Two-way ANOVA with Dunnett’s multiple comparisons test. *\*P* ≤ 0.05, \*\**P* ≤ 0.01, \*\*\**P* ≤ 0.001, \*\*\*\**P* ≤ 0.0001.

**Figure 6:**
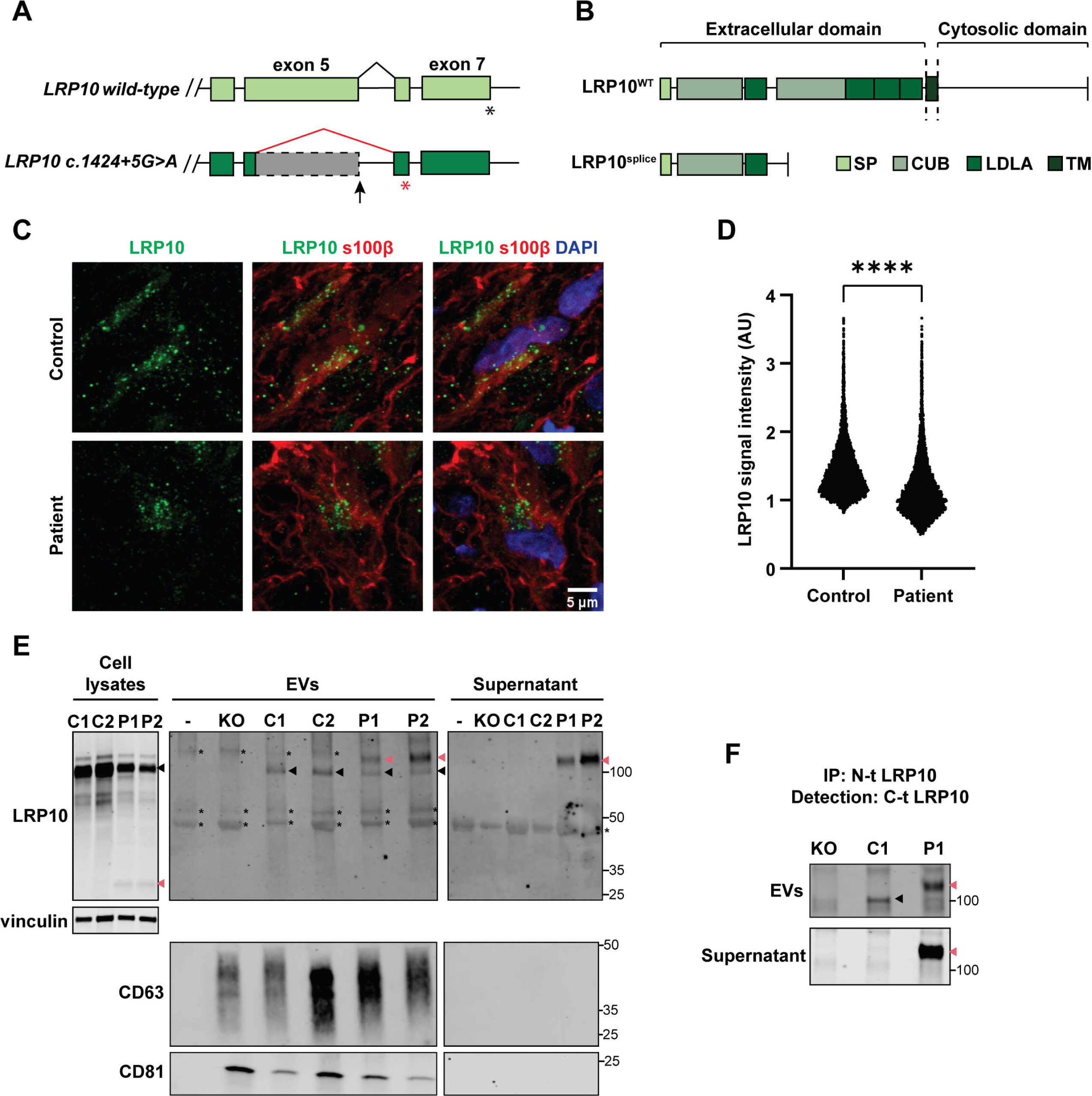
LRP10 splice variant expression and secretion in patient-derived hMLOs and astrocytes. (**A**) Schematic of the effect on splicing of the *c.1424+5G>A LRP10* variant. (**B**) Schematic of the result at the protein level of the *c.1424+5G>A LRP10* variant (LRP10^splice^). SP = signaling peptide; CUB = complement C1r/C1s, Uegf, Bmp1 domain; LDLA = low-density lipoprotein A domain; TM = transmembrane. (**C**) Representative confocal images of LRP10 in s100β-positive astrocytes in control and LRP10^splice^ patient-derived hMLOs. (**D**) Quantification of LRP10 objects signal intensity in s100β-positive astrocytes reported in log scale. N ≥ 8793 LRP10 objects. (**E**) Representative western blot of cell lysates, EVs, and supernatant fractions from LRP10^KO^, control (C1 and C2 lines), and patient iPSC-derived astrocytes (P1 and P2 clones). “-” indicates media that has not been in contact with cells. Black arrowheads indicate full-length LRP10, pink arrowheads indicate aberrant LRP10 species, and “*” indicates non-specific binding. (**F**) LRP10 pulldown from the EVs and supernatant fractions in (**E**). An antibody against the N-terminal domain (N-t) of LRP10 was used to immunoprecipitate LRP10, and an antibody against the C-terminal domain (C-t) was used for detection. All individual data points are shown. Log- transformed data were analysed by Mann Whitney test. \*\*\*\**P* ≤ 0.0001.

**Figure 7:**
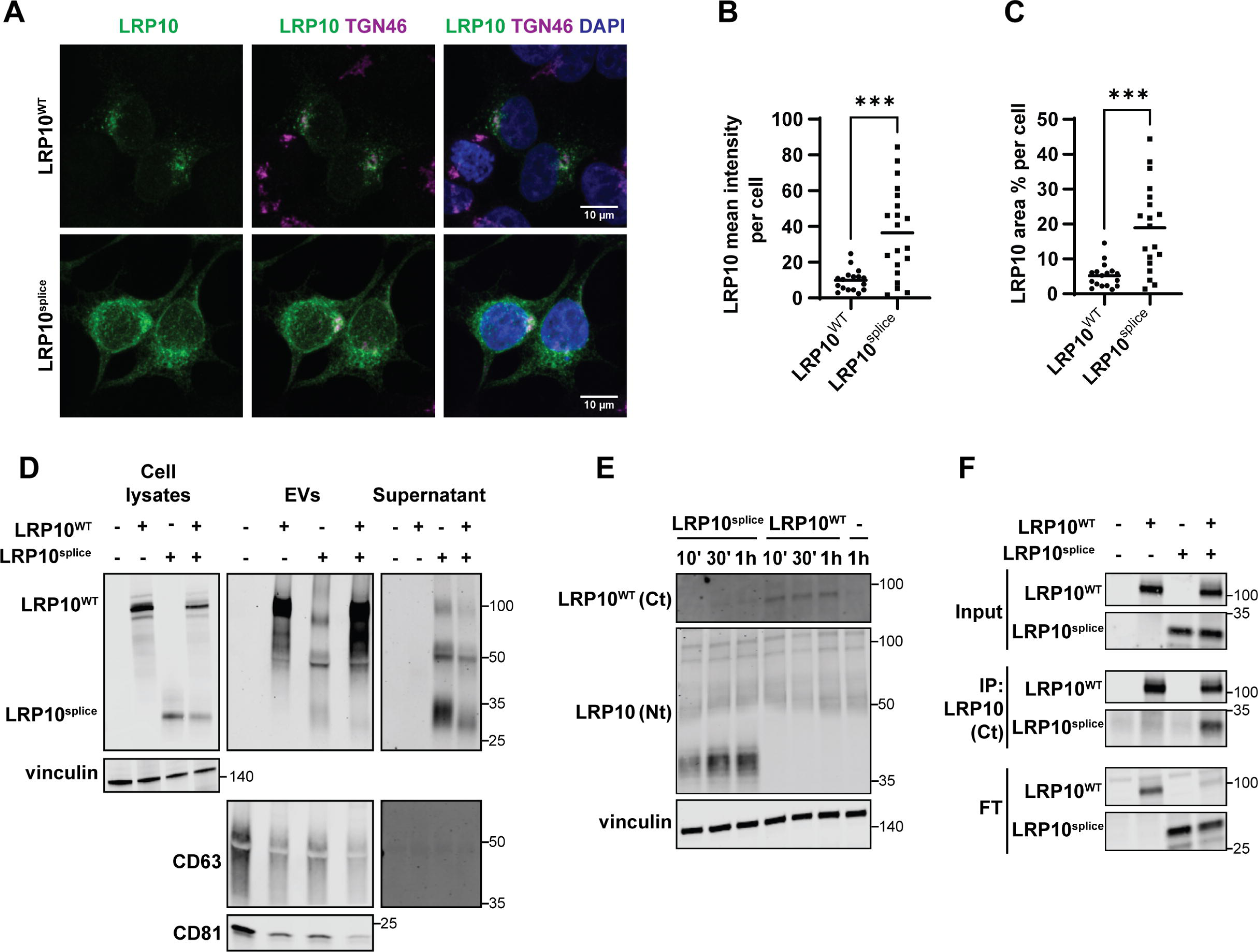
LRP10^splice^ localisation, secretion, and uptake. (**A**) Representative confocal images of LRP10^WT^ and LRP10^splice^ in overexpressing LRP10^KO^ HEK-293T cells relative to TGN46. (**B**, **C**) Quantifications of LRP10^WT^ and LRP10^splice^ mean signal intensity (**B**) and LRP10 area % (**C**) per cell. N = 17 and 19 biological replicates, respectively. (**D**) Representative western blot of cell lysates, EVs, and supernatant fractions from LRP10^splice^–expressing cells. (**E**) LRP10^WT^ and LRP10^splice^ uptake in LRP10^KO^ cells using antibodies against the C-terminal (Ct) and the N- terminal (Nt) domains of LRP10 over time. (**F**) LRP10 immunoprecipitation with the antibody against the Ct domain of LRP10 that does not bind to LRP10^splice^. All individual data points are shown including the mean. Data were analysed by unpaired T-test. \*\*\**P* ≤ 0.001.

To study the involvement of other secretory pathways in LRP10-mediated α-synuclein secretion, we focused on pathways that we demonstrated to affect intra- and extracellular LRP10 levels (Fig. 1E-G). To this aim, we used LRP10 and α-synuclein-overexpressing cells and analysed α- synuclein secretion after the different treatments. BFA treatment induced a significant increase in α-synuclein secretion over time in control cells, but not in LRP10-overexpressing cells (Fig. 5D and 5E), suggesting that BFA treatment induces an alternative α-synuclein secretion pathway only in control cells. Next, Thapsigargin treatment induced a more significant and prolonged decrease in α-synuclein secretion in LRP10-overexpressing cells (Fig. 5F and 5G). In contrast, treatment with BafA1 resulted in a decrease in α-synuclein levels in both control and LRP10-overexpressing cells (Fig. 5H and 5I). Finally, MG-132 treatment, which blocks the proteasome degradation pathway, significantly blocked LRP10-mediated α-synuclein secretion, with no effect on α- synuclein secretion in control cells (Fig. 5J and 5K). Combined, these data show that LRP10-mediated α-synuclein secretion is influenced by ER stress and proteasomal function.

### *LRP10*-variant-carrying patient cells show defects in LRP10 expression and secretion

Work from Quadri et al. (9) identified two heterozygous *LRP10* genetic variants in genomic position *c.1424+5* in DLB and PD patients. These variants were shown to interfere with *LRP10* mRNA splicing, resulting in the appearance of a premature stop codon in the variant-carrying allele (Fig. 6A and Supplementary resource 1), which ultimately results in a shorter truncated protein lacking most functional domains and the transmembrane domain (LRP10^splice^, Fig. 6B).

Here, we studied LRP10^splice^ in cells derived from a late-onset DLB patient carrying a heterozygous *c.1424+5 G→A* variant in *LRP10* (9). We first generated iPSC lines derived from this patient and a non-demented control and characterised them via immunocytochemistry and qPCR for expression of pluripotency markers (Table 1, Supplementary Fig. 4, control-3 and patient clones 1-3). hMLOs were generated from control-3 and patient clone-3 iPSCs and were characterised via immunocytochemistry (Supplementary Fig. 5A, 4 months-old hMLOs) and western blot analyses (Supplementary Fig. 5B-F, 2 and 4 months-old hMLOs), which revealed the presence of neurons (MAP2, tubulin-β3, and α-synuclein), astrocytes (GFAP), and oligodendrocytes (QKI-7) in both lines. Both the control-3 and the patient clone-3 lines showed a significant increase in GFAP levels from 2 to 4 months in culture (Supplementary Fig. 5F), similar to the previously characterised control-1 line (Supplementary Fig. 3E). We subsequently analysed LRP10 distribution in control and LRP10^splice^ hMLOs. Since astrocytes present high endogenous LRP10 levels (11, 55, 56), we analysed LRP10 expression in s100β-positive astrocytes via immunocytochemistry. As a result, we found a significant reduction in LRP10 signal intensity within s100β-positive astrocytes in patient hMLOs when compared to controls (Fig. 6C and 6D). However, western blot analyses revealed that the total amount of LRP10 in hMLOs was not significantly different between the two hMLO lines (Supplementary Fig. 5B and 5C). Therefore, LRP10^splice^ hMLOs produce relevant neural cell types and present an altered LRP10 signal distribution in s100β-positive astrocytes.

To further investigate the patient-derived LRP10 species, two iPSC clones from the LRP10^splice^ variant-carrying patient (clones 1 and 2: P1 and P2) and two control lines (controls-1 and -2: C1 and C2) iPSC lines were differentiated into astrocytes according to published protocols (11, 49).

Patient iPSCs were characterised via immunoctyochemistry and qPCR for expression of pluripotency markers (Supplementary Fig. 4, patient clones 1 and 2). To determine the efficiency of the differentiation protocol, astrocyte cultures were characterised at 2 months after the start of the differentiation via immunocytochemistry, which showed expression of the astrocytic markers GFAP, SOX9, AQP4, and s100β (Supplementary Fig. 6). After 4 months of differentiation, LRP10 intra- and extracellular distribution was investigated in astrocyte cell lysates and fractionated conditioned media via western blot (Fig. 6E). In cell lysates, the wild-type LRP10 protein was detected in both controls and patient-derived astrocytes (Fig. 6E, cell lysates, black arrowheads), and an additional ∼30 kDa LRP10 species was only detected in cell lysates from the patient P1 and P2 astrocytes (Fig. 6E, cell lysates, pink arrowheads). This observation is in line with the predicted size of the truncated protein that results from the mutant allele. Conditioned media was fractionated into EVs and supernatant fractions as described earlier (Supplementary Fig. 1A). EVs isolated from media that had not been in contact with cells (“-”) did not show the presence of CD63 and CD81 (Fig. 6E, EVs), suggesting the lack of EVs. This sample and EVs isolated from LRP10^KO^ cells were used to distinguish LRP10 signal from non-specific binding of the antibody (Fig. 6E, EVs and Supernatant, asterisks). Interestingly, although LRP10^WT^ was found in the EVs fraction of both control and patient cells (Fig. 6E, EVs, black arrowheads), an additional high molecular weight LRP10 species was detected in EVs derived from patient cells (Fig. 6E, EVs, pink arrowheads). Strikingly, this additional high molecular weight LRP10 protein could also be detected in the supernatant fraction derived from the patient cells (Fig. 6E, Supernatant, pink arrowheads). To determine the nature of the novel secreted high molecular weight LRP10 species, we immunoprecipitated LRP10 from the control and patient EVs and Supernatant fractions using an antibody against the N-terminal domain (N-t) of full-length LRP10, which binds to both LRP10^WT^ and LRP10^splice^. For detection of the immunoprecipitated proteins by western blot analysis, an antibody against the C-terminal domain (C-t) of full-length LRP10 was used, which only recognises LRP10^WT^ and not LRP10^splice^. In both the EVs and supernatant fractions from the patient line (P1) the additional high molecular weight LRP10 protein was detected with the C-t antibody, therefore confirming the presence of full-length LRP10^WT^ protein in this high molecular weight fragment (Fig. 6F, pink arrowheads). The cause of the increased LRP10 molecular weight is unknown and remains to be determined. A single protein fragment corresponding to the expected LRP10^WT^ molecular weight was observed only in the EVs fraction and not in the Supernatant of the control cell fractions (Fig. 6F, black arrowhead). In conclusion, LRP10^splice^-carrying iPSC-derived astrocytes express an additional intracellular ∼30 kDa LRP10 species, in addition to a novel secreted LRP10 form of high molecular weight in both EVs and supernatant media fractions.

### Overexpressed LRP10^splice^ is localised, secreted, and internalised aberrantly

To further characterise the patient-derived LRP10^splice^, we generated an overexpression construct consisting of the patient’s mutant cDNA sequence (9) (Supplementary resource 1). We transfected this construct and a plasmid containing LRP10^WT^ in LRP10^KO^ HEK-293T cells and studied their intracellular localisation via immunocytochemistry (Fig. 7A). LRP10^WT^ localised to vesicles that clustered around the perinuclear region, and partially colocalised with trans-Golgi network 46 kDa protein (TGN46) (9, 11–14). However, LRP10^splice^ presented a more intense (Fig. 7B) granular-like pattern distributed around the whole cell cytoplasm (Fig. 7A and 7C). Next, we investigated intra- and extracellular LRP10^splice^ distribution in cell lysates and conditioned media fractions via western blot (Fig. 7D). In cell lysates we found an additional ∼30 kDa band only in LRP10^splice^– expressing cells, which was comparable to the molecular weight of the product found in *LRP10 c.1424+5>A* patient-derived astrocytes (Fig. 6E). In conditioned media, LRP10^WT^ was exclusively detected as a ∼100 kDa band in the EVs fraction (Fig. 7D, EVs) as shown previously (Fig. 1B and Supplementary Fig. 1B). In contrast, cells expressing LRP10^splice^ or a combination of LRP10^splice^ and LRP10^WT^ secreted multiple LRP10 species ranging from 28 to 110 kDa in both EVs and supernatant fractions (Fig. 7D, EVs and Supernatant). Therefore, LRP10^splice^ presents a disrupted intracellular localisation pattern and generates an intracellular LRP10 species of similar molecular weight to the one found in patient-derived cells and is aberrantly secreted in both EVs and supernatant fractions in the form of multiple protein fragments of different molecular weights.

As we have demonstrated that LRP10 protein can be internalised via clathrin-mediated endocytosis (Fig. 2), we next investigated LRP10^splice^ uptake in LRP10^KO^ cells. Conditioned media from LRP10^WT^ or LRP10^splice^-overexpressing cells was placed on LRP10^KO^ cells for 10 min, 30 min, and 1 h before lysing the acceptor cells. We observed that LRP10^splice^ was internalised more efficiently than LRP10^WT^, which could only be visualised using an antibody against the C-terminal domain of LRP10 (Fig. 7E). Interestingly, internalised LRP10^splice^ consisted of several protein species of different molecular weights ranging from 30 to 40 kDa, and not the original 30 kDa band or the 28-110 kDa bands found in media (Fig. 7D). These observations suggest that LRP10^splice^ is internalised more efficiently than LRP10^WT^ and is processed into 30-40 kDa protein species.

Next, to study whether LRP10^splice^ physically interacts with LRP10^WT^, we immunoprecipitated LRP10 using an antibody directed against the C-terminal domain of full-length LRP10, which does not recognize LRP10^splice^. Western blot analysis revealed that LRP10^splice^ co-immunoprecipitates with LRP10^WT^, demonstrating their physical interaction (Fig. 7F). Taken together, these data indicate that the PD and DLB-associated *LRP10 c.1424+5* variants encode a truncated LRP10^splice^ protein which displays a severely disrupted secretion and internalisation patterns and can physically interact with LRP10^WT^.

### The LRP10 splice variant antagonises wild-type LRP10-mediated α-synuclein regulation

We next studied the effect of LRP10^splice^ on LRP10-mediated α-synuclein regulation. To do so, we first transfected LRP10^KO^ HEK-293T cells with *SNCA* and different amounts of LRP10^WT^ and LRP10^splice^ and inspected cell lysates and conditioned media 48 h post-transfection (Fig. 8A). As a result, we observed that LRP10^splice^ overexpression led to a decrease in LRP10^WT^ levels (Fig. 8B), and it counteracted LRP10^WT^-mediated α-synuclein intracellular levels and secretion (Fig. 8C and 8D). This effect on α-synuclein secretion was not completely caused by the decrease in LRP10^WT^ levels, since overexpression of α-synuclein and LRP10^splice^ alone in LRP10^KO^ cells led to decreased α-synuclein intracellular levels compared to control cells (Supplementary Fig. 7). Additionally, in contrast to wild-type LRP10, α-synuclein overexpression did not affect LRP10^splice^ levels (Supplementary Fig. 7B). We also studied the effect of LRP10^splice^ expression on α- synuclein levels in later time points. We generated a dox-inducible LRP10^splice^ cell line in LRP10^KO^ HEK-293T cells via antibiotic selection, and kept LRP10^splice^ expression active during the duration of the experiment. α-Synuclein was transfected and intracellular α-synuclein levels were analysed via western blot 72, 96, and 120 h after transfection (Fig. 8E). As a result, we observed that LRP10^splice^ expression led to a significant increase in α-synuclein levels 72 h after transfection (Fig. 8F), in contrast to the previously observed decrease at 48 h (Fig. 8C and Supplementary Fig. 7). Combined, these results suggest a possible dominant negative effect of the LRP10 splice variant.

**Figure 8:**
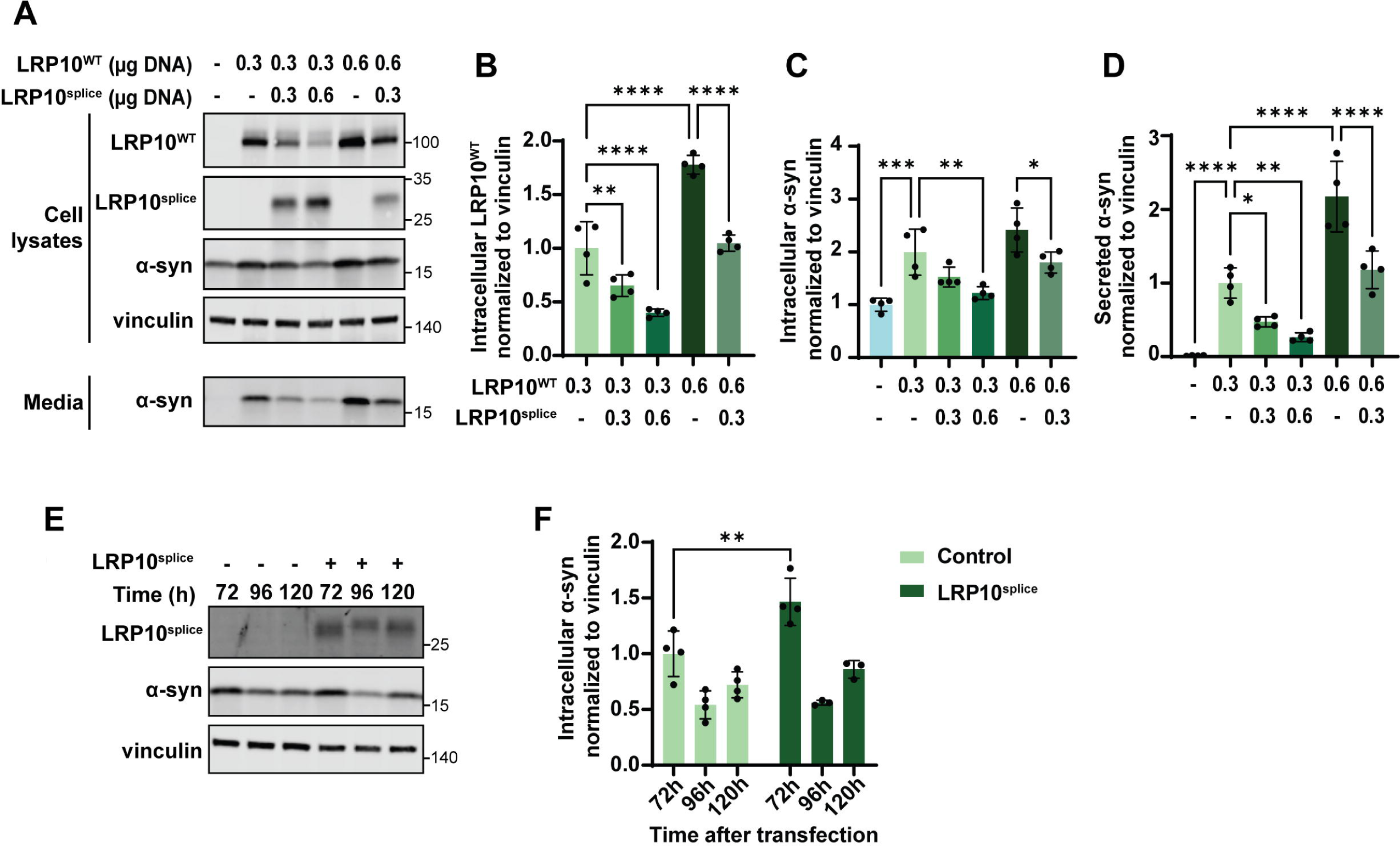
Effect of LRP10^splice^ on LRP10^WT^–mediated α-synuclein regulation. (**A**) Representative western blot of cell lysates and conditioned media from LRP10^WT^, LRP10^splice^, and α-synuclein-overexpressing cells. (**B-D**) Western blot quantifications of intracellular LRP10^WT^ (**B**) and intracellular (**C**) or extracellular α-synuclein (**D**) normalised to vinculin. N = 4 biological replicates. (**E**) Representative western blot of LRP10^splice^ and α-synuclein-overexpressing cells over time. (**F**) Quantification of intracellular α-synuclein normalised to vinculin at 72, 96, and 120 h after α-synuclein transfection. LRP10^splice^ expression was kept active via Doxycycline treatment every 2 days. N = 4 biological replicates. All data are expressed as mean ± SD with individual data points shown. Data were analysed by Two-way ANOVA with Sidak’s multiple comparisons test.

## Discussion

Our findings provide initial functional evidence for the role of LRP10 in LBDs. We first show proof for a potential mechanism that leads to the non-cell-autonomous origin of LRP10 in Lewy bodies, which is based on LRP10 secretion from astrocyte-derived EVs, and subsequent uptake and accumulation in neurons under cellular stress conditions. Furthermore, we demonstrate that LRP10 regulates α-synuclein intracellular levels and strongly induces α-synuclein secretion in an ER and proteasome-dependent manner. Additionally, we characterised the patient-derived LRP10^splice^ variant, and we show that it presents abnormal localisation, secretion, and uptake patterns. Finally, our findings indicate that the LRP10^splice^ variant behaves in a dominant negative fashion towards the regulatory role of wild-type LRP10 in controlling intra- and extracellular α- synuclein levels.

LRP10 is a distant member of the LDLR family (10). Interestingly, some members of this family, namely LDLR, LRP1, LRP1B, LRP4, and SORL1, have been identified as key players in different neurodegenerative diseases (65–71). Particularly, LRP1 was recently identified to regulate the transmission of tau and α-synuclein in human neurons *in vitro* and in mice (66–68). On another note, LRP10 has been shown to physically interact with SORL1, which suggests that they might function together in a cellular context. Given the fact that both LRP10 and LRP1 have strong effects on α-synuclein secretion, it is therefore possible that LRP1 and LRP10 have coordinated roles in α-synuclein transmission, perhaps through physical interaction, for which additional research is required.

LRP10 was found in significant amounts in the core of Lewy bodies in LBD patients, even though it was almost undetectable in control neurons (11). This observation raised the question of whether, under pathological conditions, LRP10 is secreted from surrounding cells and taken up and accumulated in neurons. In this work, we show that LRP10 is efficiently secreted from overexpressing HEK-293T cells and control astrocytes via EVs (Fig. 1A-D, 1J, 6E, 7D, and Supplementary Fig. 1B). EVs are essential in transcellular material transfer and cellular communication, for instance, between astrocytes and neurons (72). Under pathological conditions, EVs contents have been shown to change (73), and they have even been suggested to promote disease spread (64, 74–76). Since the current work focuses on LRP10 secretion *in vitro*, follow-up studies with patient brain material will be essential to clarify whether LRP10 dosage and structure differ in EVs from LBD patients when compared to controls, and if it may influence disease progression. In addition to generating valuable knowledge about disease pathogenesis, it would represent a potential biomarker, since EVs are easily detected in blood and cerebrospinal fluid (CSF) (77).

In addition to being linked to EVs, we show that LRP10 secretion is promoted after inducing autophagic dysfunction (Fig. 1E-L). Particularly, BafA1-treated astrocytes secrete higher levels of LRP10 in whole media but also in EVs (Fig. 1J-L). It has been previously described that inhibition of the autophagy-lysosome degradation pathway can trigger protein secretion through EVs as a compensatory mechanism to dispose of cellular material, including α-synuclein (63, 78, 79). Therefore, LRP10 secretion via EVs possibly becomes exacerbated when its degradation via the lysosome-autophagy pathway is impaired via similar mechanisms. In this work, we also show that BafA1 treatment in hMLOs induces the formation of LRP10-positive objects in neurons (Fig. 3F and 3G). Since LRP10 expression in neurons is very low and the treatment duration was only of 4 h, we believe the origin of these neuronal LRP10 structures not only consists of an accumulation of endogenous LRP10. Alternatively, BafA1 treatment might induce LRP10 secretion from nearby cells (e.g., astrocytes), which is subsequently internalised by these neurons. Given that defects in autophagy are very well described in LBDs (62), it is possible that pre-existing defects in degradation pathways or other cellular stresses trigger LRP10 accumulation in Lewy bodies in both idiopathic and genetic LBD cases. We also hypothesise that, in patients carrying pathogenic *LRP10* variants, the initial trigger for cellular stress might be the presence of abnormal LRP10 species. However, it is yet to be determined whether wild-type or mutant LRP10 play an active role in Lewy body formation and maturation or if LRP10 localisation in Lewy bodies is a secondary effect of the disease.

In healthy brains and hMLOs, LRP10 expression is higher in astrocytes than in neurons (11, 55–57) (Figure 3A and 3B). A growing amount of evidence supports the essential role of astrocytes in LBD pathogenesis. Astrocytes are central in keeping brain homeostasis and perform essential functions such as the secretion of survival factors and synapse regulation in neurons and maintaining the blood–brain barrier (80–83). In PD and other neurodegenerative disorders, astrocytes have been proven inefficient in their neuronal-supporting role, in addition to inducing neuroinflammation (84–86). Besides *LRP10*, several other LBD genes, including *DJ-1*, *PINK1*, and *GBA*, are highly expressed in astrocytes, and the encoding proteins have been shown to have a role in astrocyte biology (55, 87–91). Moreover, disease-linked genetic variants have been shown to interfere with their normal function and contribute to disease progression (87, 91–93). Additional research will clarify the role of LRP10 in astrocytic function, and whether it is impaired in *LRP10*-variant-carrying patients.

Our findings show that LRP10 overexpression leads to time-dependent changes in α-synuclein intracellular levels (Fig. 4). Additionally, we show that α-synuclein overexpression leads to a decrease in LRP10 levels (Fig. 4A and 4B). Similar feedback loops have been described before between α-synuclein and LBD-linked proteins, such as GBA (94). α-Synuclein is considered to be central in LBDs, and a key defining feature of Lewy bodies, but whether and how α-synuclein initiates the disease cascade in all patients remains unclear. Moreover, even though the majority of LBD research has focused on the detrimental effect of α-synuclein, several arguments challenge this hypothesis. For instance, triple synuclein KO mice present neuropathological defects and reduced survival (95, 96). Therefore, proteins that regulate α-synuclein levels may provide relevant insight into α-synuclein biology and its role in disease initiation and progression. Furthermore, even though a myriad of α-synuclein species has been described, clear evidence for their significance in pathology is lacking for the majority of them (32). In contrast to α-synuclein oligomers, which have a well-documented neurotoxic effect, this work focuses on monomeric α- synuclein, with a poorly understood pathogenic role (32, 97). However, several modifications on α-synuclein monomers are associated with increased misfolding and aggregation potential (32, 34, 98). Additionally, increased *SNCA* dosage leads to a strong neuropathological phenotype, also in humans carrying gene multiplications (99, 100). Therefore, although we show a clear link between LRP10 and monomeric α-synuclein, further research comparing monomeric and other forms of α-synuclein will provide useful insight to understand whether this interaction is protective or detrimental to disease initiation and progression.

In this study, we show that LRP10 overexpression also induces a strong increase in α-synuclein secretion (Fig. 4 and 5). Interestingly, LRP12, another distant relative of the LDLR family, was also found to modify α-synuclein cell-to-cell transfer *in vitro* (101). α-Synuclein transmission is currently considered to be central in disease pathogenesis and progression (29). Oligomeric α-synuclein is detected in CSF and it has been proposed as a biomarker for PD (102). Therefore, LRP10-mediated α-synuclein secretion might have diagnostic potential for *LRP10*-variant-carrying patients, although it is not yet clear if these patients present a loss or gain of LRP10 function. α-Synuclein has been shown to be secreted via several unconventional pathways (63, 64, 78, 79). Here, we demonstrate that LRP10 induces α-synuclein secretion via cell-stress response pathways, including ER stress and the proteasome (Fig. 5F, 5G, 5J, and 5K). In line with this observation, several studies show that α-synuclein aggregates or overexpression lead to defects in proteasomal function (103–105). Interestingly, certain cellular sensors that are able to detect misfolded proteins have been shown to induce proteasome-dependent secretion of these proteins (106). Taking this into account, it is possible that LRP10 is acting as a sensor of α-synuclein overexpression and exporting misfolded proteins through a novel unconventional α-synuclein secretion pathway.

The present work focuses on studying the overexpression of LRP10 and α-synuclein simultaneously in the same cells. As previously discussed, LRP10 is enriched in astrocytes in comparison to neurons, whereas α-synuclein is typically considered to be a neuronal protein. To explain the interaction between LRP10 and α-synuclein, a multicellular model is required. One possible explanation is that LRP10 regulates α-synuclein in a later disease state within neurons after LRP10 internalisation. Another possibility is that LRP10 regulates α-synuclein processing within astrocytes, which, in LRP10-variants-carrying patients, might play a role in disease initiation. Increasing evidence suggests that astrocytes play a more important role in α-synuclein pathology than what was initially considered. Recent work from Altay et al. (107) described that astrocytic α-synuclein accumulations present unique post-translational modifications, and colocalise with different markers when compared to neuronal α-synuclein-positive inclusions. For this reason, on many occasions, astrocytic α-synuclein is not detected with antibodies that recognize α-synuclein in neuronal Lewy bodies. In agreement, several reports show different types of α-synuclein accumulations in astrocytes in patient post-mortem material (108–117). Furthermore, pathological α-synuclein has been shown to dysregulate essential functions in astrocytes (87, 92, 118–120), and, in turn, astrocytes have been shown to spread α-synuclein pathology to neurons (84, 121). Taken together, the present study demonstrates a functional link between LRP10 and α-synuclein, but the biological context where this interaction takes place possibly involves several cell types and remains of difficult interpretation.

Finally, we show several defects associated with the LRP10 splice variant identified in PD and DLB patients (Fig. 6-8). First, we observe differences in LRP10 localisation in LRP10^splice^ hMLOs, since LRP10 signal is reduced within s100β-positive astrocytes as determined by immunocytochemistry (Fig. 6C and 6D), but not in total protein lysates from hMLOs as determined by WB (Supplementary Fig. 5B and 5C). Next, we show that patient-derived cells secrete an aberrant patient-specific high molecular weight LRP10 species (Fig. 6E). Interestingly, this high molecular weight form of LRP10 is not found intracellularly, pointing towards a patient-specific mechanism occurring during secretion. Crucially, we show that this patient-specific extracellular LRP10 species contains wild-type LRP10 (Fig. 6F), but it is not found in control cells. Given the size of this aberrant extracellular LRP10 form on SDS-page gels (Fig. 6E and 6F), one possibility might be that LRP10^splice^ forms SDS-resistant protein oligomers with LRP10^WT^. In line with this hypothesis, it has been described that LRP6 forms SDS-resistant homodimers via its LDLA domain to be functionally active as a co-receptor for Wnt signalling (122). It is therefore possible that LRP10 also dimerises via its LDLA domains, which become SDS-resistant due to the molecular properties of the LRP10^splice^ variant. Alternatively, LRP10^splice^ might increase LRP10^WT^ molecular weight by mediating its binding to other proteins or the introduction of post-translational modifications. Furthermore, the fact that LRP10^WT^ is found in the supernatant fraction of the media from patient cells suggests that it is likely not linked to cell membranes, which is striking considering LRP10 is a type-I transmembrane protein (10). These results suggest that the LRP10^splice^ variant also affects LRP10^WT^ processing, either via proteolytic processing at the plasma membrane or other mechanisms that can dissociate LRP10 from the membrane.

In our overexpression model, LRP10^splice^ shows a different intracellular localisation pattern when compared to LRP10^WT^ (Fig. 7A-C). In contrast to LRP10^WT^, which presents a vesicular localisation that accumulates predominantly around the nucleus, LRP10^splice^ presents a more intense cytoplasmic distribution with a granular pattern. Furthermore, LRP10^splice^ is secreted in the form of different protein species of higher molecular weight (Fig. 7D). However, extracellular overexpressed LRP10^splice^ cDNA presents a different pattern on SDS-PAGE gels when compared to endogenous expression in the *c.1424+5G>A LRP10* variant-carrier patient lines (Fig. 6E), suggesting that the *LRP10* genetic context in the genome, the LRP10^WT^ - LRP10^splice^ ratio, the expression time, and/or the cell type might play a role in this secretion pattern. Additionally, LRP10^splice^ is found as several protein species of different sizes after uptake, and not the original monomeric 30 kDa form (Fig. 7E). These findings suggest that secreted LRP10 in patients carrying the *c.1424+5G>A* variant might have aggregation potential; an effect that is perhaps not occurring in the donor cells (e.g. astrocytes), but might occur in the acceptor cells (e.g., neurons). Furthermore, we demonstrate that LRP10^splice^ binds to full-length LRP10 (Fig. 7F) and antagonises its function in α-synuclein regulation (Fig. 8A-D), pointing to a dominant-negative effect. Additionally, overexpression of LRP10^splice^ alone in LRP10^KO^ cells affects α-synuclein levels in an opposite way when compared to LRP10^WT^ (Fig. 8E and 8F, and Supplementary Fig. 7). Therefore, not only LRP10^splice^ has a dominant negative effect over LRP10^WT^ but might also regulate α-synuclein levels. Overall, we show evidence for the pathological potential of the LRP10 splice variant, which might be disease-causing in these carriers. Additional work remains ahead to further elucidate the consequences of this variant in mutation-carrying patients. For instance, neuropathological examination of these patients remains to be performed. Furthermore, since several different types of *LRP10* variants have been identified (9, 16–25), they might affect LRP10 function in different ways, which remains to be investigated.

In conclusion, this work provides initial evidence for the non-cell autonomous origin of LRP10 in Lewy bodies, in addition to a functional role of LRP10 in LBDs via α-synuclein regulation, and shows pathogenic mechanisms linked to the patient-associated *c.1424+5G>A LRP10* variant.

## Availability of data and materials

The data that support the findings of this study are available from the corresponding authors, upon reasonable request.

## Supporting information

Supplementary statistics

## Abbreviations

AA: Ascorbic acid
AD: Alzheimer’s disease
APP: Amyloid precursor protein
AQP4: Aquaporin 4
AU: Arbitrary Units
BafA1: Bafilomycin A1
BDNF: brain-derived neurotrophic factor
BFA: Brefeldin A
BSA: Bovine serum albumin
CNTF: Ciliary neurotrophic factor
CQ: Chloroquine
CSF: Cerebrospinal fluid
C-t: C-terminal
dbcAMP: Dibutyryl cyclic adenosine monophosphate
DLB: Dementia with Lewy bodies
DMEM: Dulbecco’s modified eagle medium
DMSO: Dimethylsulfoxid
DTT: dithiothreitol
EGF: Epidermal growth factor
ER: Endoplasmic Reticulum
EVs: Extracellular vesicles
FBS: Fetal bovine serum
FGF: Fibroblast growth factor
GBA: Glucosylceramidase
GDNF: Glial cell line-derived neurotrophic factor
GFAP: Glial fibrillary acidic protein
gRNA: Guide RNA
HEK-293T: Human embryonic kidney 293T
hMLOs: human iPSC-derived midbrain-like organoids
HUES: Human embryonic stem cell lines
ICC: Immunocytochemistry
iPSCs: Induced pluripotent stem cells
IRES: Internal ribosome entry site
KO: Knock-out
LBDs: Lewy body diseases
LDLR: Low-density lipoprotein receptor
LRP10: Low-density lipoprotein receptor-related protein 10
NeoR: Neomycin resistance gene
NPCs: Neural precursor cells
N-t: N-terminal
NTA: Nanoparticle tracking analysis
PAGE: Polyacrylamide gel electrophoresis
PBS: Phosphate buffered saline
PD: Parkinson’s disease
PDD: Parkinson’s disease dementia
PFA: Paraformaldehyde
PuroR: Puromycin resistance gene
QKI-7: Quakin 7
RT-qPCR: Reverse transcription quantitative real-time PCR
SAG: Smoothened Agonist
SDS: Sodium dodecylsulfate
SORL1: Sortilin related receptor 1
TBS: Tris buffered saline
TEM: Transmission electron microscopy
TGF-β3: Transforming growth factor beta 3
TH: Tyrosine hydroxylase
WB: Western blot
WT: Wild-type

## Acknowledgements

We thank the donors and their families for making this study possible. We also thank Erasmus MC iPS Core Facility for providing the hiPSC lines.

## Author information

### Authors and Affiliations

**Department of Clinical Genetics, Erasmus Medical Center, Rotterdam, The Netherlands**

Ana Carreras Mascaro, Martyna M. Grochowska, Valerie Boumeester, Ece Naz Bilgiҫ, Guido J. Breedveld, Vincenzo Bonifati & Wim Mandemakers.

**Department of Neurology, Erasmus Medical Center, Rotterdam, The Netherlands**

Leonie Vergouw & Frank Jan de Jong.

**Department of Urology, Erasmus Medical Center, Rotterdam, The Netherlands**

Natasja F. J. Dits.

**Department of Pathology, Erasmus Medical Center, Rotterdam, The Netherlands**

Martin E. van Royen.

#### Contributions

ACM, MMG, MEvR, and WM designed the experiments. ACM, MMG, VaB, ENB, GJB, NFJD, and MEvR performed the experiments. ACM, MMG, VaB, NFJD, and MEvR performed data analysis. ACM, MMG, VaB, NFJD, MEvR, VB, and WM interpreted the data. LV and FJdJ provided the patient biopsy. ACM wrote the initial draft of the manuscript. VB and WM conceived the study and supervised the experiments. All authors critically read the manuscript and approved the final version.

### Ethics approval and consent to participate

Primary cell lines were obtained within study protocols approved by the medical ethical committee of Erasmus MC and conformed to the principles of the Declaration of Helsinki. The participating subjects provided written informed consent for the use of the material for research purposes.

