## Supplementary material for "LRP10 as a novel α-synuclein regulator in Lewy body diseases"

ATGCTGTTGGCCACCCTCCTCCTCCTCCTTGGAGGCGCTCTGGCCCATCCAGA
CCGGATTATTTTTCCAAATCATGCTTGTGAGGACCCCCCAGCAGTGCTCTTAGAA
GTGCAGGGCACCTTACAGAGGCCCCTGGTCCGGGACAGCCGCACCTCCCCTGCCA
ACTGCACCTGGCTCATCCTGGGCAGCAAGGAACAGACTGTCACCATCAGGTTCCA
GAAGCTACACCTGGCCTGTGGCTCAGAGCGCTTAACCCTACGCTCCCCTCTCCAG
CCACTGATCTCCCTGTGTGAGGCACCTCCCAGCCCTCTGCAGCTGCCCGGGGGCA
ACGTCACCATCACTTACAGCTATGCTGGGGCCAGAGCACCCATGGGCCAGGGCTT
CCTGCTCTCCTACAGCCAAGATTGGCTGATGTGCCTGCAGGAAGAGTTTCAGTGC
CTGAACCACCGCTGTGTATCTGCTGTCCAGCGCTGTGATGGGGTTGATGCCTGTG
GCGATGGCTCTGATGAAGCAGCATCTTTGCCCCCCTCTCCCGGATGGAGGCTGAG
ATTGTGCAGCAGCAGGCACCCCCTTCCTACGGGCAGCTCATTGCCCAGGGTGCCA
TCCCACCTGTAG

**Supplementary resource 1**: **sequence encoding for LRP10$^{splice}$.**

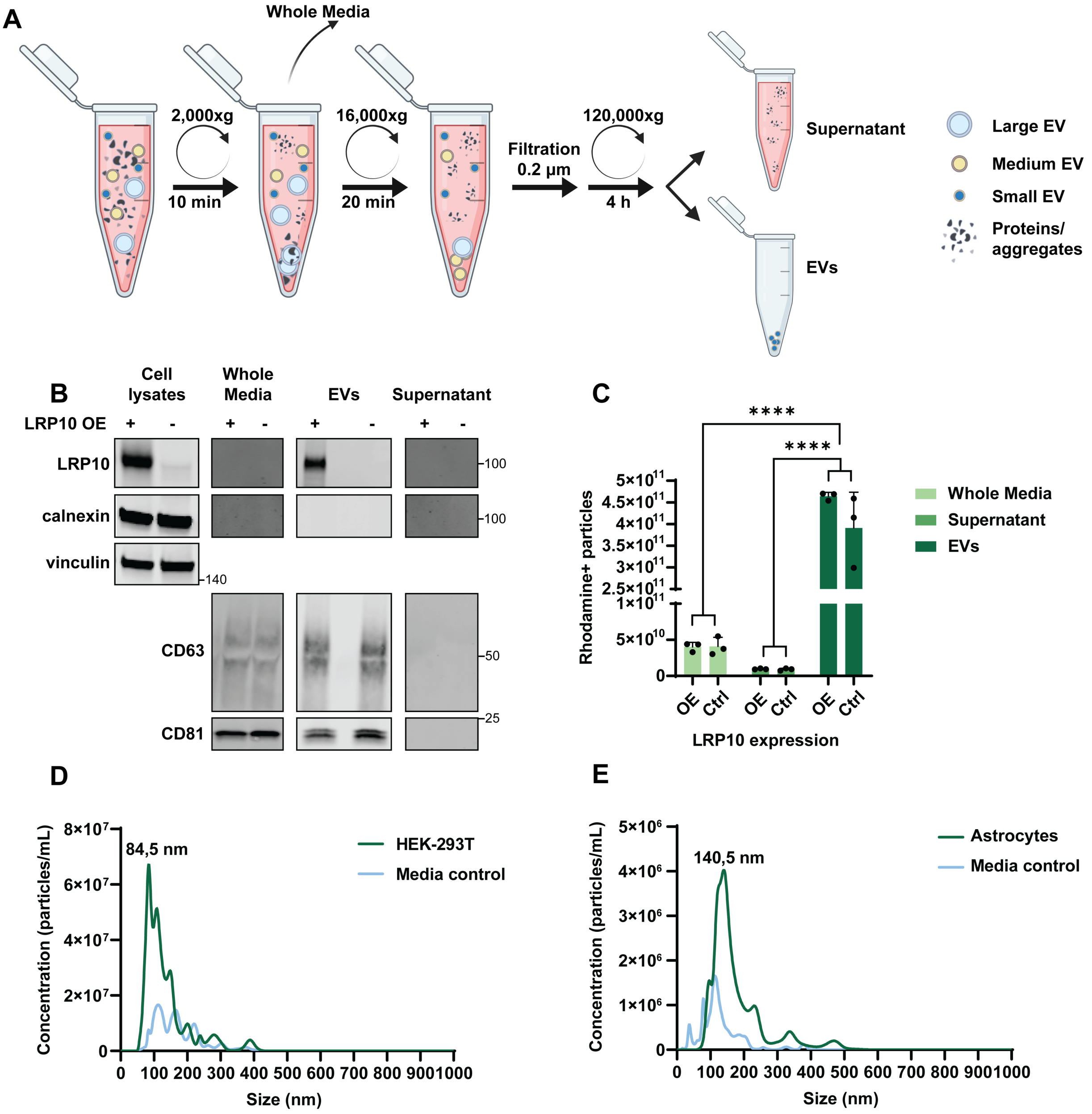


**Supplementary Figure 1: Extracellular vesicles (EVs) isolation protocol and quality control.** (**A**) Schematic of the ultracentrifugation and filtration protocol for EVs isolation from conditioned media. (**B**) Representative western blot probed for calnexin, vinculin, CD63, and CD81 in different media fractions from LRP10-overexpressing (OE) and control (Ctrl) HEK-293T cells. (**C**) EVQuant of Rhodamine-positive particles in different fractions resulting from the EVs isolation protocol. N = 3 biological replicates. (**D-E**) Nanoparticle tracking analysis showing concentration and diameter sizes (nm) of isolated EVs from HEK-293T cells (**D**), iPSC-derived astrocytes (**E**), or control media that has not been in contact with cultured cells. All data are expressed as mean ± SD with individual data points shown in (**C**). Data were analysed by Two-way ANOVA with Sidak's multiple comparisons test. ****$P \leq 0.0001$.

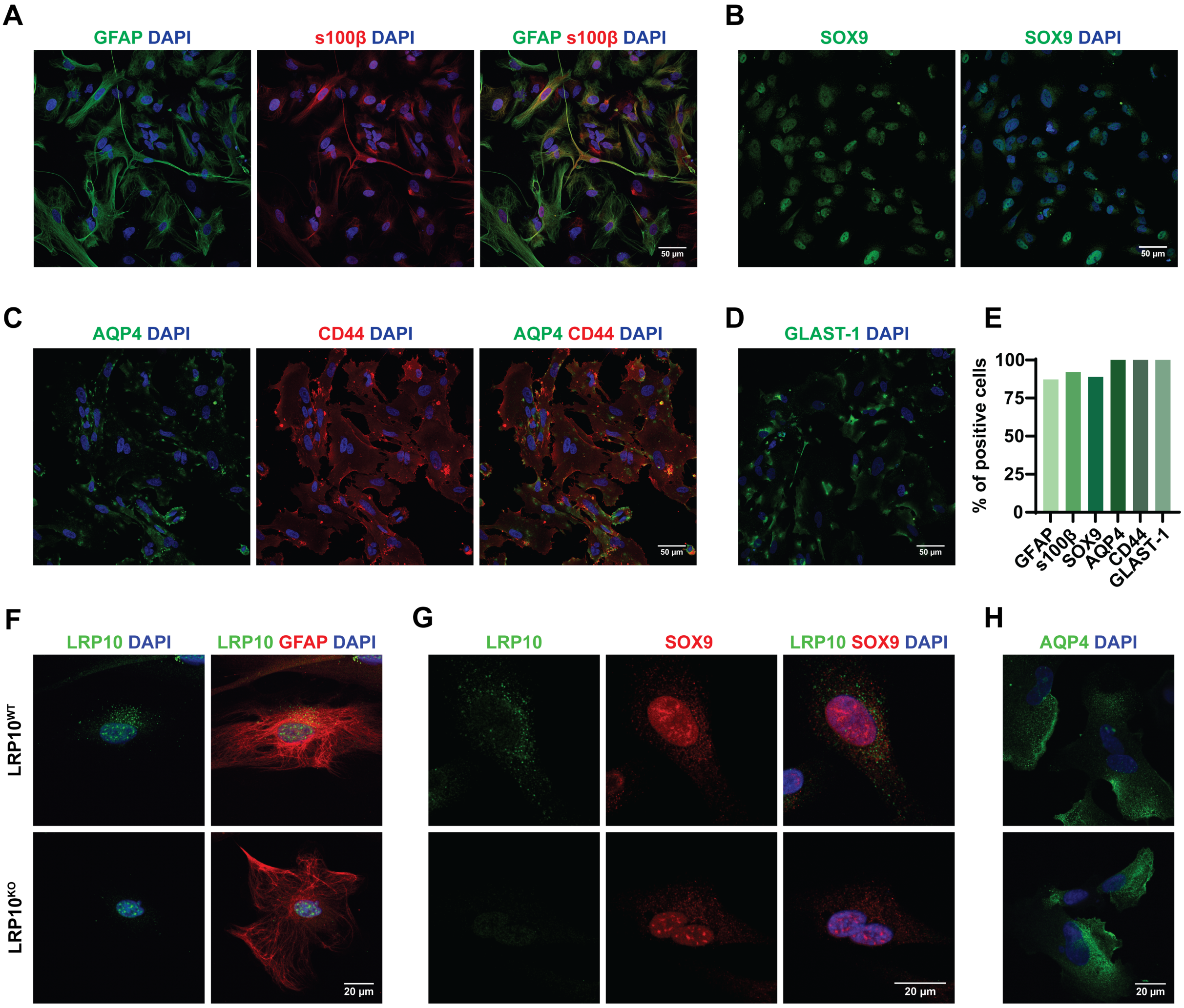


**Supplementary Figure 2: Characterisation of control and LRP10$^{KO}$ iPSC-derived astrocytes.** (**A-D**) Representative confocal images of 4 months-old control-2 iPSC-derived astrocytes stained for the astrocytic markers GFAP and s100β (**A**), SOX9 (**B**), AQP4 and CD44 (**C**), and GLAST-1 (**D**). (**E**) Quantification of % of positive cells for the specified markers from (**A-D**). (**F-H**) Characterisation of wild-type LRP10 (LRP10$^{WT}$) and LRP10 KO (LRP10$^{KO}$) astrocytes, immunostained for the presence of LRP10 together with GFAP (**F**), LRP10 together with SOX9 (**G**), and AQP4 (**H**). The antibody raised against the C-terminal domain of LRP10 was used in (**F**).

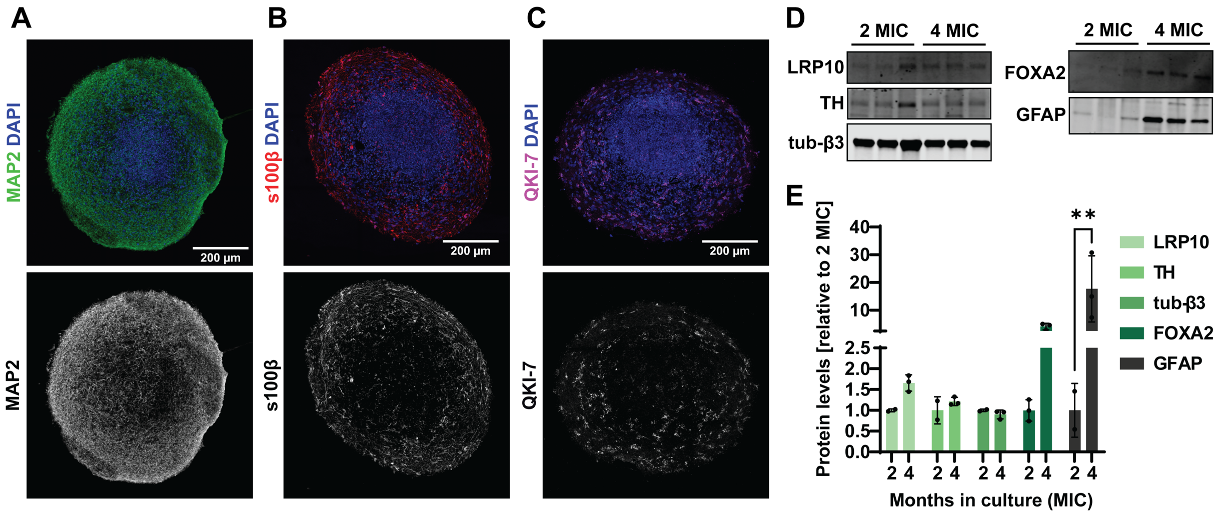


**Supplementary Figure 3: Characterisation of control hMLOs.** (**A-C**) Representative tile scans of 2 months-old control-1 hMLOs showing MAP2 (**A**), s100β (**B**), and QKI-7 (**C**) immunostainings. (**D**) Representative western blot of 2 and 4 months-old (months in culture, MIC) control-1 hMLOs. Blots were probed for expression of LRP10, TH, tubulin β3 (tub-β3), FOXA2 and GFAP. (**E**) Western blot quantification of the indicated markers relative to 2 MIC. N = 3 hMLOs per condition. All data are expressed as mean ± SD with individual data points shown. Data were analysed by Two-way ANOVA with Sidak's multiple comparisons test. $^{**}P \leq 0.01$.

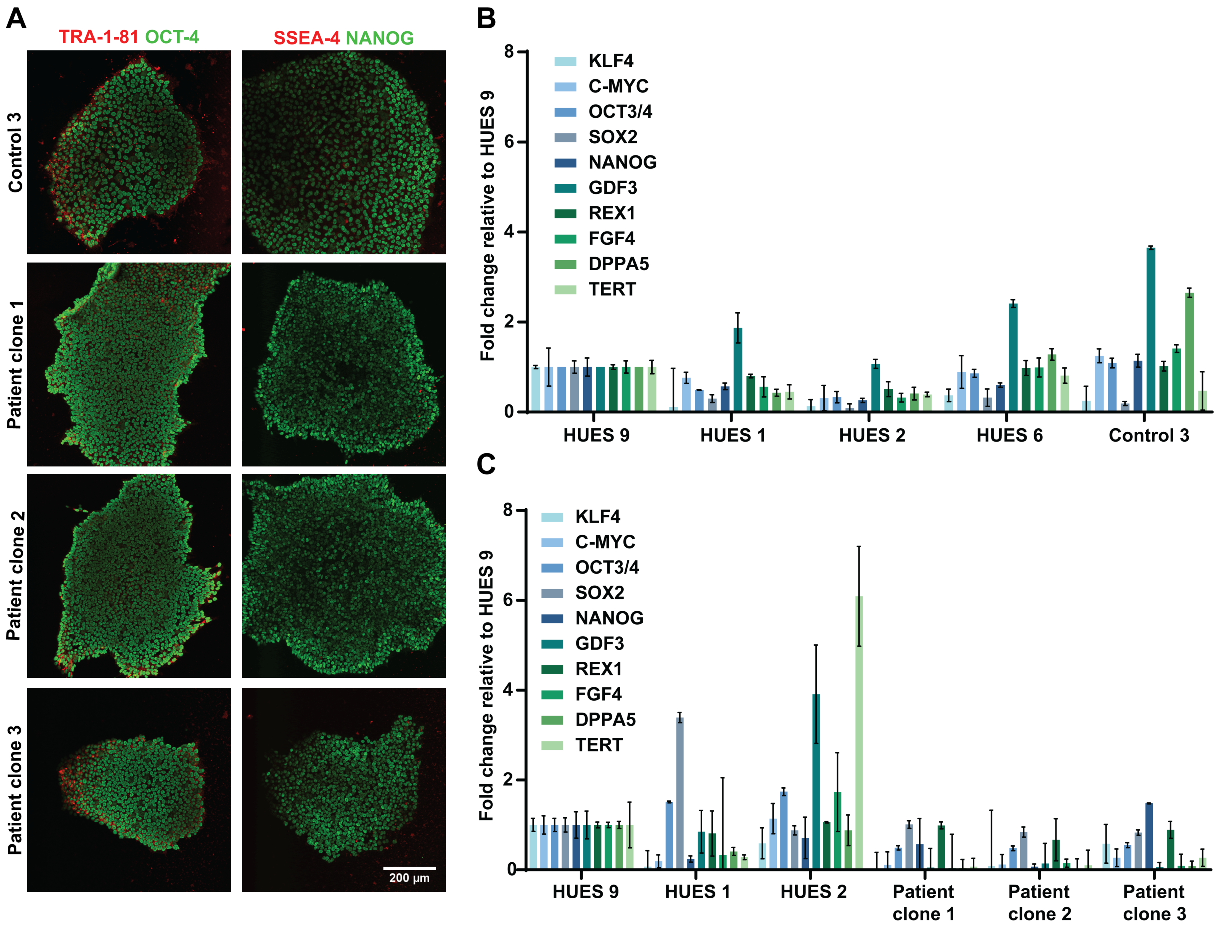


**Supplementary Figure 4: Characterisation of iPSC lines.** (**A**) Representative confocal images of pluripotency markers OCT4, TRA-1-81, NANOG, and SSEA-4 in control-3 and three iPSC clones from the *c.1424+5G>A LRP10*-variant carrying patient. (**B**) qPCR for pluripotency markers in control-3 in comparison to control lines HUES 9, 1, and 2. N = 3 technical replicates. (**C**) qPCR for pluripotency markers in the 3 clones derived from the *LRP10*-variant carrying patient in comparison to control lines HUES 9, 1, and 2. N = 3 technical replicates. All data are expressed as mean ± SD.

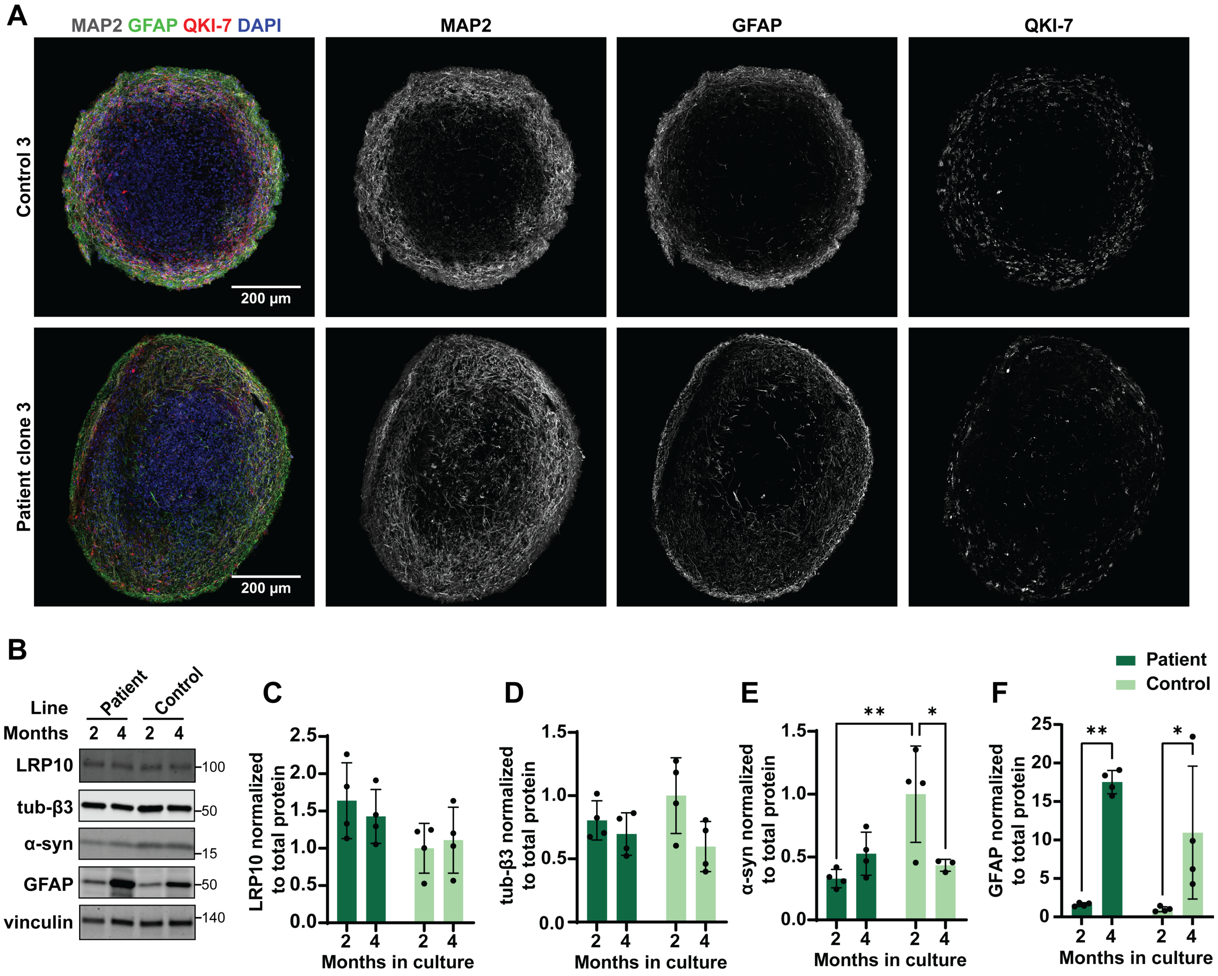


**Supplementary Figure 5: Characterisation of control-3 and LRP10$^{splice}$ patient clone-3 hMLOs.** (**A**) Representative tile scans of 4 months-old control-3 and patient clone-3 hMLOs showing the distribution of MAP2, GFAP, and QKI-7-positive cells. (**B**) Representative western blot of 2 and 4 months-old control and patient hMLOs probed fo expression of LRP10, tubulin β3, α-synuclein, GFAP, and vinculin. (**C-F**) Quantifications of LRP10 (**C**), tubulin β3 (**D**), α-synuclein (**E**), and GFAP (**F**) normalised to total protein. N = 4 biological replicates. All data are expressed as mean ± SD with individual data points shown. Data were analysed by Two-way ANOVA with Sidak's multiple comparisons test. *$P \leq 0.05$, **$P \leq 0.01$.

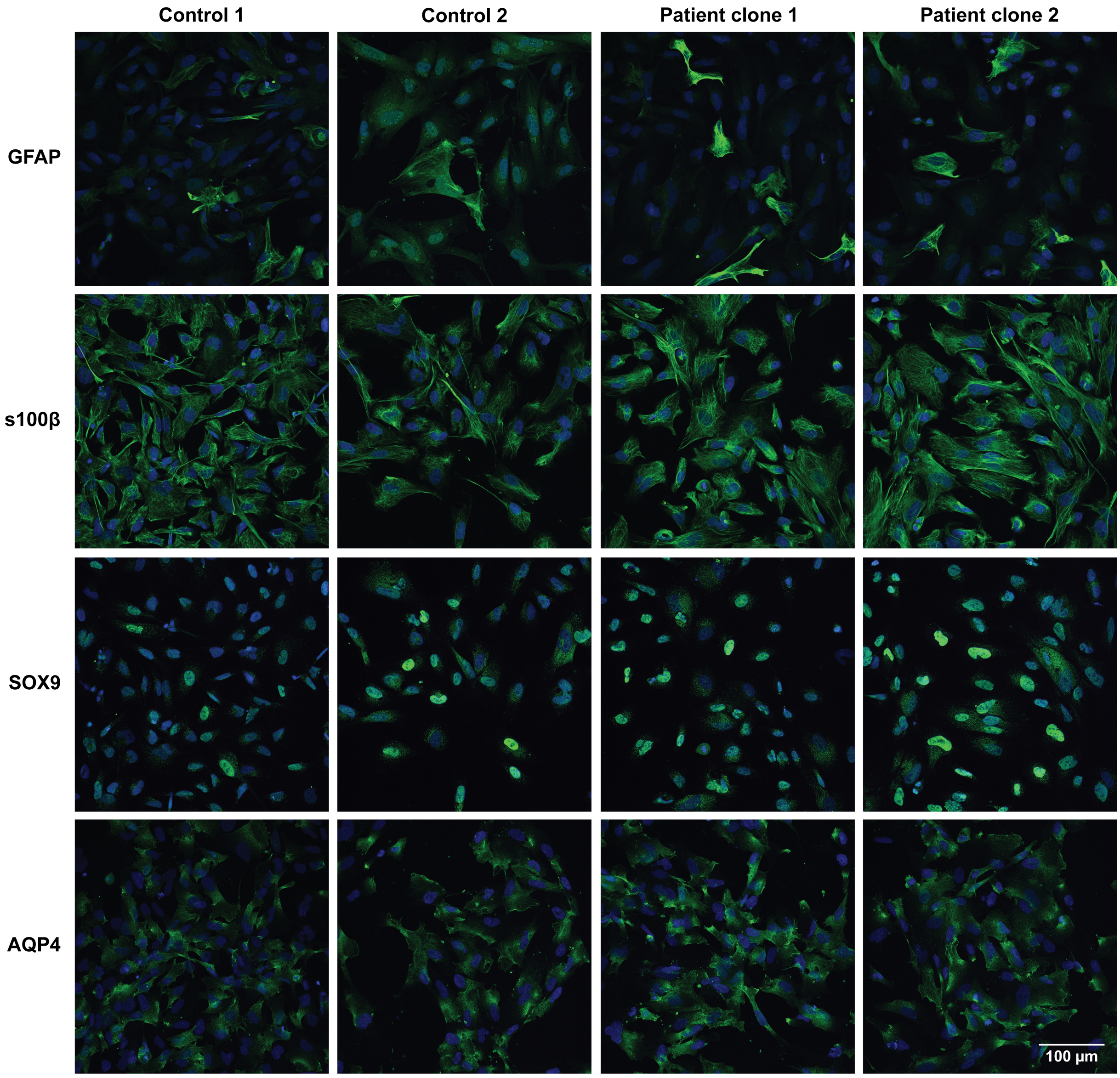


**Supplementary Figure 6: Characterisation of iPSC-derived astrocyte lines.** Representative confocal images of the astrocytic markers GFAP, s100β, SOX9 and AQP4 (in green) and DAPI (in blue) in 2 months-old iPSC-derived astrocytes from control lines 1 and 2, and patient clones 1 and 2.

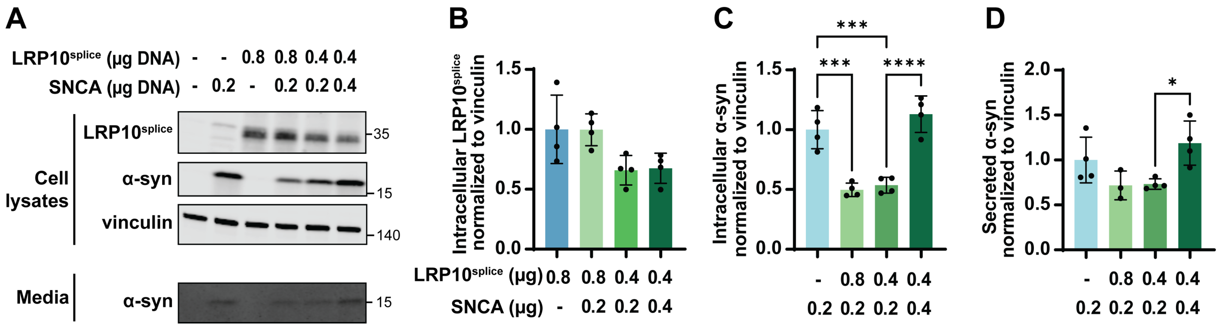


**Supplementary Figure 7: LRP10$^{splice}$ and α-synuclein overexpression in HEK-293T cells.** (**A**) Representative western blot of cell lysates and conditioned media from HEK-293T overexpressing LRP10$^{splice}$ and α-synuclein. (**B-D**) Western blot quantifications of intracellular LRP10$^{splice}$ (**B**), intracellular α-synuclein (**C**), and secreted α-synuclein (**D**) normalised to intracellular vinculin. N = 4 biological replicates. All data are expressed as mean ± SD with individual data points shown. Data were analysed by One-way ANOVA with Tukey's multiple comparisons test. $*P \leq 0.05$, $***P \leq 0.001$, $****P \leq 0.0001$.
